# Environmental activity-based protein profiling for function-driven enzyme discovery from natural communities

**DOI:** 10.1101/2022.11.11.516116

**Authors:** Sabrina Ninck, Thomas Klaus, Tatiana V. Kochetkova, Sarah P. Esser, Leonard Sewald, Farnusch Kaschani, Christopher Bräsen, Alexander J. Probst, Ilya V. Kublanov, Bettina Siebers, Markus Kaiser

## Abstract

Microbial communities are significant drivers of global biogeochemical cycles, yet accurate function prediction of their proteome and discerning their activity *in situ* for bioprospecting remains challenging. Here, we present environmental activity-based protein profiling (eABPP) as a novel proteomics-based approach bridging the gap between environmental genomics, correct function annotation and *in situ* enzyme activity. As a showcase, we report the successful identification of active thermostable serine hydrolases by combining genome-resolved metagenomics and mass spectrometry-based eABPP of natural microbial communities from two independent hot springs in Kamchatka, Russia. eABPP does not only advance current methodological approaches by providing evidence for enzyme and microbial activity *in situ* but also represents an alternative approach to sequence homology-guided biocatalyst discovery from environmental ecosystems.

## Introduction

Microbial organisms account for the vast majority of the unexplored natural biodiversity^1^ and colonize all of the earth’s conceivable ecological niches, thereby forming microbial communities of distinct complexity and fluctuating composition^2,3^. Many of the terrestrial and marine environmental microbial communities thriving under moderate to extreme conditions in diverse ecosystems such as soil, freshwater, seawater or hot springs harbor unique genes with (novel) associated functionalities^2^. Their set of enzymes equips them with diverse metabolic activities, some of which yet unknown, that turn these communities into a promising resource for environmental biotechnology^4^. Especially thermostable enzymes that can function as biocatalysts in diverse biotechnological processes are sought with intense demand^5^ and have fostered systematic bioprospecting of hydrothermal microbial communities^6–9^. Their metagenomic-based investigation has turned into a feasible biocatalysts discovery approach, in particular for screening organisms or communities that are non-culturable using standard techniques^10,11^. The subsequent functional analysis is commonly done by sequence-driven bioinformatic prediction of protein function^12^. This approach, however, is hampered by a large quantity of proteins of unknown function (i.e hypotheticals) or misannotated enzymes as well as the presence of large protein superfamilies, for which functional predictions are still often insufficient^13^. ‘Functional metagenomics’ approaches can help to overcome these limitations by complementing sequence-based approaches with an activity-based screening after construction of a metagenomic library^14^. They enable the discovery of novel enzyme classes but require elaborate and challenging cloning and expression efforts with subsequent biochemical characterization^12,15^. Moreover, this approach delivers no information on the expression and, as applies to all ‘omics’-based strategies, the activity state of an enzyme-of-interest in its natural environment.

Since its first application more than two decades ago, activity-based protein profiling (ABPP) has emerged into a powerful approach for studying enzyme activities. Within this period, a huge variety of activity-based probes (ABPs) that target different enzymes or even whole enzyme classes have been developed^16–18^. A ‘classical’ ABP is composed of a reactive warhead, a tag and, if present, a chemical or peptidic linker region. While the warhead is often an electrophile-containing inhibitor molecule that targets a nucleophile at the active site of an enzyme for covalent labeling, the tag is used for target detection or enrichment^19^. Thus, ABPP not only allows the visualization of labeled enzymes via the use of fluorescent reporter groups, but also enables target enzyme identification by mass spectrometry (MS) if a moiety for target enrichment, such as biotin, is used as a reporter group^20^. Accordingly, an appropriate use of enzyme or enzyme class-specific ABPs enables the functional annotation of enzymes^21^. With the integration of ‘click chemistry’ into the ABPP workflow, two-step chemical probes have emerged, which facilitate a simple *in vivo* application of ABPP under physiological conditions^22–24^.

In the last years, ABPP was mainly used in the context of biomarker or drug discovery as well as *in vivo* imaging^25^; beyond that, diverse applications in micro- or plant biology, including the study of pathogens or host-pathogen interactions, have been frequently reported^26,27^. Only recently, the use of ABPP in biocatalyst discovery for industrial applications, for example for elucidating lignocellulose-degrading enzymes, has emerged^28^. In addition, first reports on the use of ABPP for studying microbial communities, with a focus on host-associated microbial communities, have been made^29^. Mayers *et al*., for example, combined ABPP with a cysteine-reactive probe and quantitative metaproteomics to identify enzymes with an altered expression in inflammatory bowel disease in a mouse model^30^. For the identification of host as well as microbiome proteins from murine fecal samples, a Comprehensive Protein Identification Library (ComPIL) database constructed from publicly available genome data^31^ was used. The exploration of environmental microbial communities by ABPP, in contrast, still relies mostly on profiling isolated strains of environmental microbes rather than on the direct profiling of complex communities^32–36^. Recently, an activity-based labeling of ammonia- and alkane-oxidizing bacteria from complex microbial communities using 1,7-octadiyne was reported^37^. This study additionally used metagenomic sequencing for further confirmation of the functional potential of the targeted microorganisms; a generic workflow for ABPP of microbial communities in the environment providing a function-based target identification of the labeled enzymes via downstream massspectrometry is however still lacking today.

We herein describe an ABPP approach that allows the functional identification of active enzymes from complex environmental microbial communities (hereafter referred to as environmental ABPP, eABPP). Our approach is based on a combination of ABPP with genome-resolved metagenomics, facilitating the assignment of dedicated activities to enzymes from microorganisms in the environment, even to those that belong to so far uncultured or unknown microorganisms. This has become possible by a tailored sample preparation and data analysis procedure that allows to detect single active enzymes within a complex metaproteome. As a ‘proof-of-concept’, we employed an alkyne-tagged version of the well-established fluorophosphonate-based (FP) chemical probe^38^, which has been previously shown to allow ABPP of serine hydrolases even under extreme experimental conditions such as elevated temperatures or low pH^36^. Moreover, members of the serine hydrolase superfamily are widely distributed across all domains of life^39^ and are biocatalysts with broad application in biotechnological processes^40–45^. Accordingly, we profiled serine hydrolase activities of microbial communities from two different hot springs located at the Uzon Caldera (Kamchatka Peninsula, Russia) under native conditions directly at the site of sampling. For further sample processing, target enzyme enrichment and MS-based analysis, the *in vivo-* labeled sediments were transferred to the laboratory. By metagenome sequencing of an untreated reference sample, an *in silico* metaproteome was constructed that was used as a reference database to identify microbial enzymes with serine hydrolase activity.

## Results

### Workflow design

For establishing a function-based enzyme identification approach for the direct profiling of an environmental microbial community, we designed a general workflow based on a combination of ABPP with metagenome sequencing (Fig. 1). This workflow starts with the collection of a microbial community sample with sufficient biomass, here a hot spring sediment. This sample is then split into seven aliquots and six of them are used for *in vivo* labeling employing a specific two-step ABP aiming to target enzymes with the desired activity among the microbial enzyme repertoire directly at the site of sampling (eABPP). While three aliquots are treated with the respective probe, the other three replicates serve as solvent controls. The remaining sample aliquot, on the other hand, is used for extraction of eDNA. The eDNA is subsequently submitted to metagenome sequencing, enabling the generation of the microbial community-specific metaproteome database via sequence assembly, genome binning and gene prediction (Supplementary Fig. 1). In parallel, proteins from the eABPP samples are extracted and cleaned up by phenol extraction and ammonium acetate precipitation, followed by an *in vitro* ‘click’ reaction with downstream affinity enrichment of labeled enzymes. Enzyme identification is then achieved by on-bead digestion of captured enzymes with subsequent LC-MS/MS analysis using the self-assembled metaproteome database for identification of active target enzymes. Finally, enzyme function can be confirmed by recombinant expression of target enzymes and biochemical enzyme characterization.

**Figure 1.**
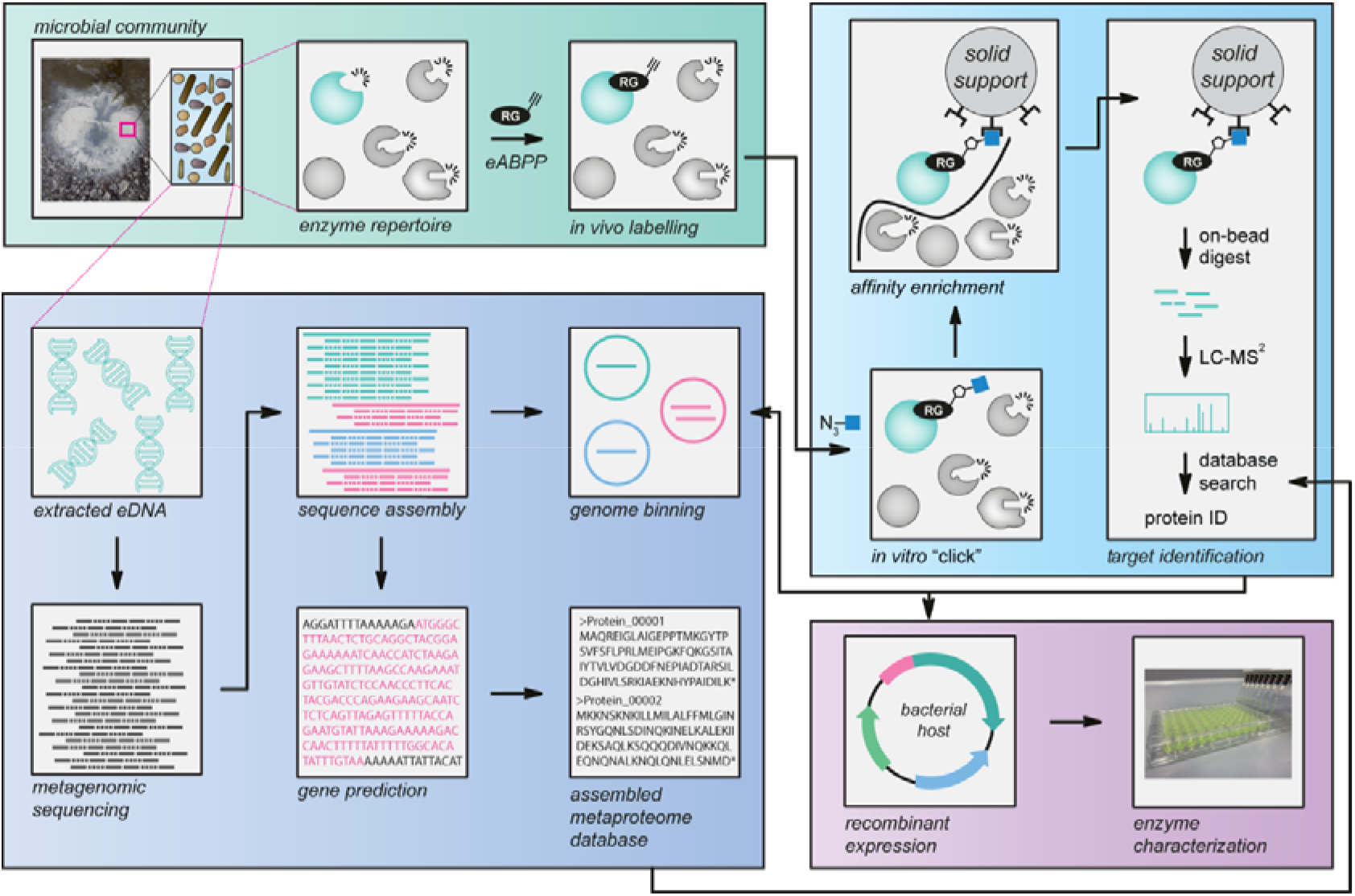
Environmental ABPP workflow. Workflow of the established environmental ABPP (eABPP) approach for the function-based identification of serine hydrolases. This approach can be divided into four different blocks, i.e. sampling and *in vivo* labeling of an environmental microbial community, metagenomics, target protein identification by LC-MS/MS and enzyme characterization of a protein of interest.

### Micerobial composition of the sampled springs

To test this eABPP approach, we performed an *in vivo* ABPP of serine hydrolases from an environmental microbial community using a FP-based ABP. Accordingly, two hot springs located in the Uzon Caldera (Kamchatka Peninsula, Russia), the ‘Arkashin shurf’ (KAM3811) and the ‘Helicopter spring’ (KAM3808), were sampled (Fig. 2). These two sites were chosen due to their physicochemical properties (a temperature in the mesothermal range (40-75°C)^46^ and a slightly acidic pH) favoring the development of an abundant microbial community. While KAM3811 is an originally artificial but for more than 30 years stable thermal pool^47^, for which 16 metagenome-assembled genomes (MAGs) have been published recently^48^, KAM3808 is a larger natural spring with no metagenome data available so far. After sampling both sites, metagenome sequencing of the epi-sedimentary microbial community was performed. We first analyzed the community composition of both sediments at the domain and class level based on the relative abundance of ribosomal protein S3 (rpS3) genes within the metagenomic datasets (relative abundance was determined via metagenomic read mapping; Fig. 3a). Both springs displayed considerable differences in terms of the percent distribution between Bacteria and Archaea and the number of different classes present within each domain. For KAM3811, 66% of the assigned species belonged to the bacterial domain, with the Aquificota being the dominating phylum, while the archaeal domain was mainly represented by the Thermoproteota (Supplementary Tab. 1). In the phenotypically and phylogenetically more diverse spring KAM3808, bacterial species account for 57 % of all organisms, with the Caldisericota making up the largest fraction, followed by the Desulfobacterota and the Aquificota. In contrast to KAM3811, archaeal species comprised mainly Euryarchaeota, while Archaea from the DPANN group made up the smallest fraction (Supplementary Tab. 2).

**Figure 2.**
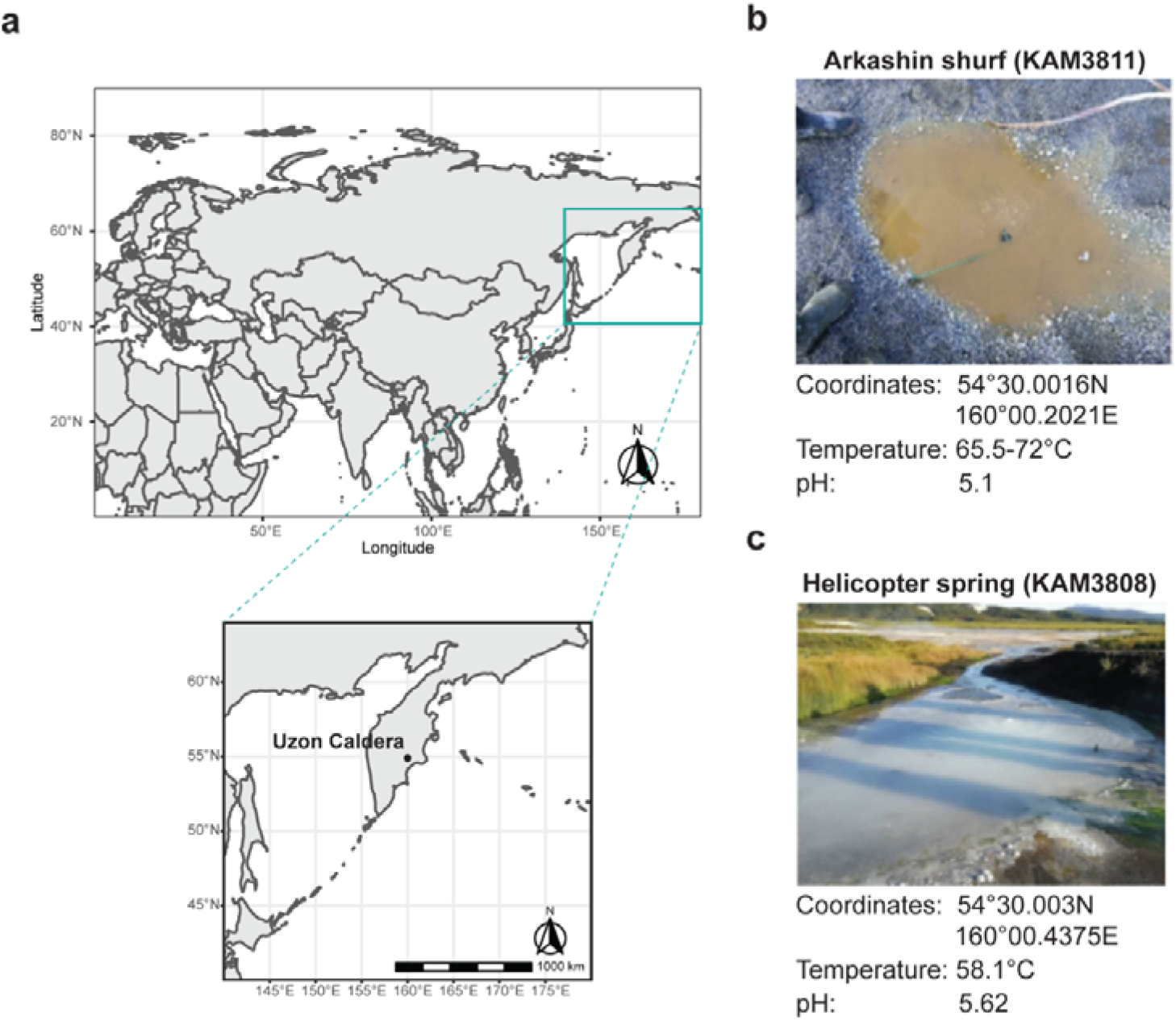
Location and community composition of the sampled springs. (**a**) The map shows the location of the two sampled springs ‘Arkashin shurf’ (KAM3811) and ‘Helicopter spring’ (KAM3808) at the Uzon Caldera region, Kamchatka Peninsula, Russia. (**b,c**) A representative picture is shown along with the exact coordinates and physicochemical properties of both springs.

**Figure 3.**
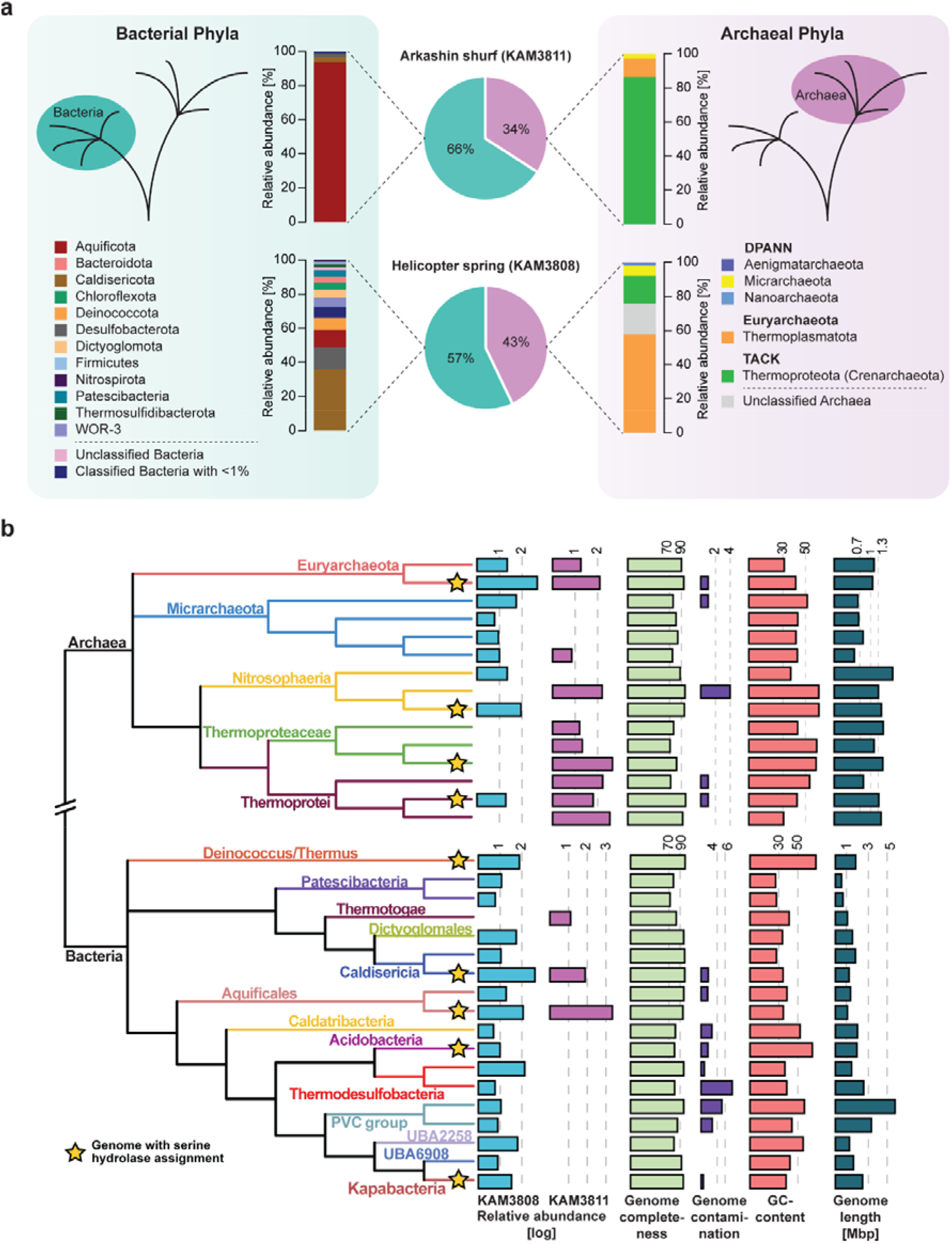
Distribution of microorganisms across KAM3811 and KAM3808. (**a**) The proportion of Bacteria (cyan) and Archaea (magenta) within the sediments sampled for eABPP is depicted as pie diagrams for KAM3811 and KAM3808. An overview of the relative distribution of representative phyla from these domains is given respectively based on the GTDB taxonomy. (**b**) Phylogenetic tree displaying the relationship between the microorganisms found across both springs as calculated with GTDB-Tk based on the dereplicated, binned and curated metagenomes from KAM3808 and KAM3811. Relative abundances of microorganisms based on the coverage of genomes are given for KAM3808 (light blue) and KAM3811 (magenta), respectively, along with their genome completeness (light green), contamination (purple), GC content (light red) and genome length (dark cyan) as calculated via checkM. The yellow stars indicate from which genomes predicted serine hydrolases were confidently identified with the applied eABPP approach.

In a more detailed analysis of these metagenomic datasets, again by relying on genome coverages, all phyla identified within the respective spring were displayed in a domain-specific phylogenetic tree with their corresponding abundancies (Fig. 3b; Supplementary File 1). For KAM3808, which displayed more heteromorphic sediments, rpS3 genes of 49 microorganisms could be identified, with *Aciduliprofundum* sp. (relative abundance of 488.1), *Caldisericum exile* (408.2) and *Caldimicrobium thiodismutans* (139.9) as the most abundant species (Supplementary Tab. 2). KAM3811, a distinctly smaller thermal pool with homogenic brown to red sediments, comprised 19 different organisms as determined from rpS3 gene analysis. Among them, *Sulfurihydrogenibium* sp. (2329.1) was identified as the dominating genus, followed by *Pyrobaculum ferrireducens* (411.3) and *Caldisphaera* sp. (389.6; Supplementary Tab. 1).

### Term-based annotation of serine hydrolases in the metaproteome database is diverse

Based on protein prediction of the assembled metagenomes, metaproteome databases consisting of 45,649 (KAM3811) and 99,930 (KAM3808) protein coding sequences, respectively, were constructed. Annotation of the metaproteome databases against the UNIREF100 database resulted in 35,323 genes encoding proteins with a predicted function for the KAM3811 dataset while 8,575 proteins remained uncharacterized. Analogously, 85,393 genes encoded proteins with a predicted function for the KAM3808 dataset, whereas 17,923 genes encoded uncharacterized proteins. The discrepancy between the number of proteins with predicted function and hypothetical ones stemmed from the lack of represented similar genes within public databases or from fragmentation of genes. Within the proteome dataset of KAM3811 and KAM3808, only two and sixteen proteins, respectively, were annotated with the term ‘serine hydrolase’. Please note that the variety of term-based annotations of enzymes within both metaproteomes that potentially belong to the serine hydrolase superfamily is huge. Predicting the actual number of serine hydrolase sequences present in the two datasets thus remains intricate, since it would require elaborate manual curation of relevant search strings. This difficulty, atop of hypothetical or misannotated proteins, represents another drawback of simple sequence-driven bioinformatic annotation when searching for dedicated enzyme activities, especially from large superfamilies, which is also addressed by our function-based eABPP approach.

### Identification of active serine hydrolases by eABPP

To identify active serine hydrolases from the sampled environmental communities, the self-constructed metaproteome databases were then employed as a reference database for the analysis of eABPP MS data resulting from an affinity enrichment of FP-alkyne-labeled target proteins via a biotin handle, on-bead digest and subsequent LC-MS/MS analysis (Supplementary File 2). Overall, 811 and 1489 protein groups were identified for the KAM3811 and KAM3808 datasets, respectively, excluding hits from the implemented contaminants database. After filtering of the initial data (see the methods section), a total of 413 protein groups for KAM3811 and 689 protein groups for KAM3808 remained for further analysis. For KAM3811, 221 protein groups displayed a positive log2-fold change when comparing the group of FP-labeled samples against the DMSO-treated control group (Fig. 4a). Of them, 15 protein groups were predicted as serine hydrolases. Most notably, these protein groups were predominantly found among the top enriched hits, with 12 protein groups found among the top 30 enriched hits (Tab. 1). For KAM3808, which showed a larger microbial diversity, 348 protein groups were found to be enriched in the group of FP-labeled samples when compared against the DMSO-treated control group (Fig. 4b). These include 31 predicted serine hydrolases. As already observed for KAM3811, most of them were top hits, with 17 protein groups found among the top 30 enriched hits (Tab. 2). To confidently predict serine hydrolase activity across the proteins that were enriched with FP-alkyne, we here complemented the protein annotation from the UniRef100 database with sequence searches against the Pfam, NCBI CCD and InterPro databases. Where necessary, additional searches against the SWISS-MODEL Template Library or with HHpred were performed to identify structural homologs (Supplementary File 3).

**Figure 4.**
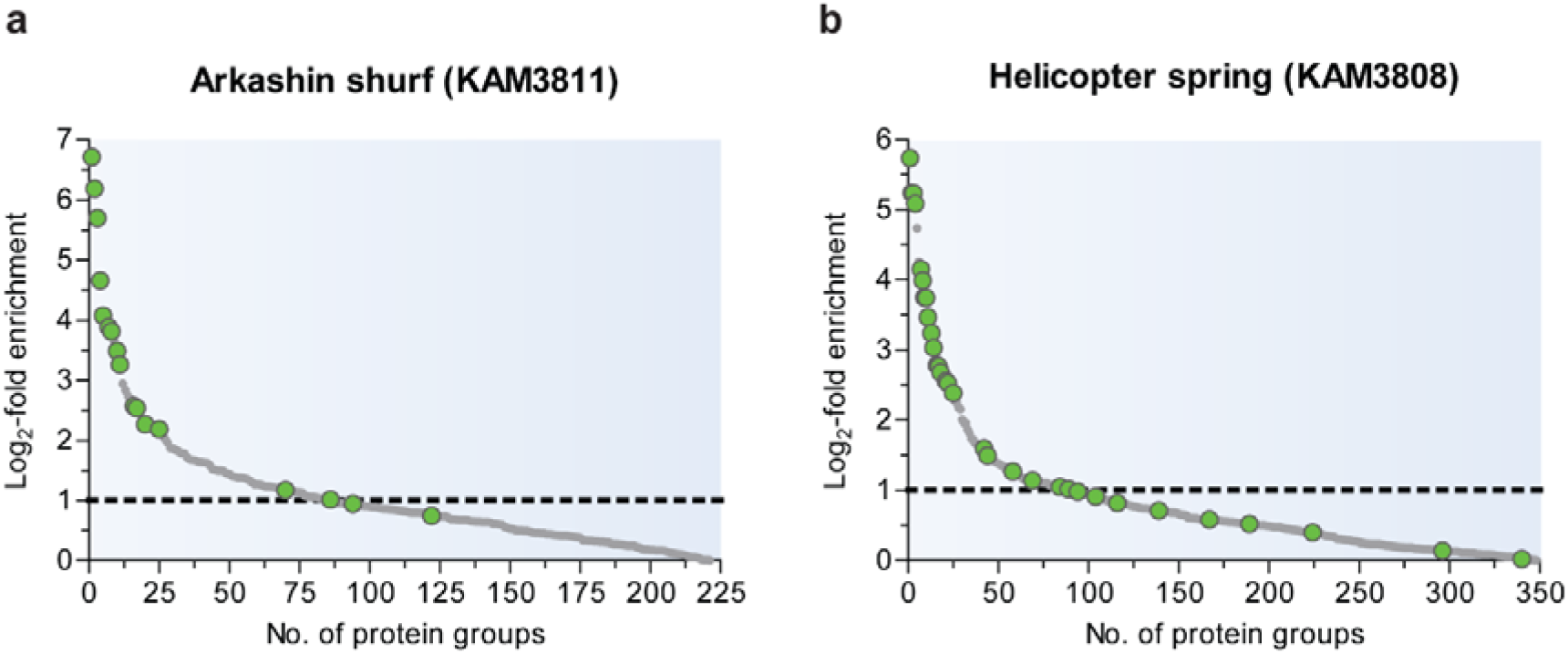
Predicted serine hydrolases identified from the sampled hot springs. Log2-fold enrichment of identified proteins labeled with FP-alkyne compared to the DMSO control for the sediments sampled from KAM3808 (**a**) and KAM3811 (**b**). Proteins predicted as serine hydrolases are displayed as green dots. Hits lying above the dotted line were more than two-fold enriched with the probe and were therefore considered primary hits. Each treatment group comprised three biological replicates.

**Table 1.** Predicted serine hydrolases identified for KAM3811.

| No. | Log2-fold<br>change | Identifier | Protein annotation<sup>#</sup> |
| --- | --- | --- | --- |
| 1 | 6.711 | ExploCarb_3811S_S4_2994_length_2665_cov_56_1 | Peptidase S8, subtilisin-related |
| 2 | 6.188 | ExploCarb_3811S_S4_103_length_35923_cov_48_7 | Penicillin/GL-7-ACA/AHL acylase |
| 3 | 5.695 | ExploCarb_3811S_S4_37_length_59942_cov_23_2 | Peptidase S8, subtilisin-related |
| 4 | 4.656 | ExploCarb_3811S_S4_179_length_24712_cov_181_27 | Penicillin/GL-7-ACA/AHL acylase |
| 5 | 4.074 | ExploCarb_3811S_S4_782_length_9153_cov_146_4 | Penicillin/GL-7-ACA/AHL acylase |
| 7 | 3.881 | ExploCarb_3811S_S4_477_length_13181_cov_9_10 | Peptidase S8, subtilisin-related |
| 8 | 3.810 | ExploCarb_3811S_S4_412_length_14764_cov_159_5 | Peptidase S8, subtilisin-related |
| 10 | 3.483 | ExploCarb_3811S_S4_5916_length_1495_cov_4_1 | Peptidase_S8/S53_dom |
| 11 | 3.258 | ExploCarb_3811S_S4_7165_length_1270_cov_6_1 | Peptidase S8, subtilisin-related |
| 17 | 2.533 | ExploCarb_3811S_S4_1591_length_4725_cov_42_1 | Protein of unknown function DUF915,<br>hydrolase-like |
| 20 | 2.264 | ExploCarb_3811S_S4_483_length_13114_cov_941_5 | Uncharacterised protein family<br>UPF0227/Esterase YqiA |
| 25 | 2.184 | ExploCarb_3811S_S4_1380_length_5439_cov_180_2 | Peptidase_S9 |
| 70 | 1.172 | ExploCarb_3811S_S4_259_length_20079_cov_121_4 | Peptidase_S49 |
| 94 | 0.943 | ExploCarb_3811S_S4_1740_length_4311_cov_157_3 | Xaa-Pro-like_dom |
| 122 | 0.745 | ExploCarb_3811S_S4_1428_length_5265_cov_777_6 | Peptidase_S49 |

<sup>#</sup> InterProScan annotation on either protein family or domain level

**Table 2.** Predicted serine hydrolases identified for KAM3808.

| No. | Log2-fold<br>change | Identifier | Protein annotation<sup>#</sup> |
| --- | --- | --- | --- |
| 1 | 5.739 | ExploCarb_3808S_S2_15464_length_1283_cov_2_1 | GDSL lipase/esterase |
| 2 | 5.244 | ExploCarb_3808S_S2_5917_length_3034_cov_1_3 | Peptidase S8, subtilisin-related |
| 3 | 5.238 | ExploCarb_3808S_S2_19633_length_1026_cov_55_1 | Putative S8A family peptidase<sup>†</sup> |
| 4 | 5.086 | ExploCarb_3808S_S2_470_length_27453_cov_4_11 | SGNH_hydro |
| 7 | 4.152 | ExploCarb_3808S_S2_593_length_22580_cov_176_21 | Esterase/lipase |
| 8 | 3.990 | ExploCarb_3808S_S2_12785_length_1520_cov_45_2 | Peptidase S8, subtilisin-related |
| 9 | 3.754 | ExploCarb_3808S_S2_10394_length_1832_cov_227_1 | Peptidase_S8/S53_dom_sf |
| 10 | 3.748 | ExploCarb_3808S_S2_1348_length_11661_cov_169_4 | PNPLA_dom |
| 11 | 3.467 | ExploCarb_3808S_S2_277_length_40468_cov_33_33 | Peptidase S8A, fervidolysin-like |
| 13 | 3.243 | ExploCarb_3808S_S2_1901_length_8595_cov_3_5 | Penicillin/GL-7-ACA/AHL acylase |
| 14 | 3.030 | ExploCarb_3808S_S2_12072_length_1604_cov_127_1 | Peptidase_S8/S53_dom_sf |
| 16 | 2.780 | ExploCarb_3808S_S2_5180_length_3427_cov_2_3 | Peptidase S8, subtilisin-related |
| 17 | 2.761 | ExploCarb_3808S_S2_12016_length_1611_cov_2_1 | AB_hydrolase_1 |
| 18 | 2.684 | ExploCarb_3808S_S2_2184_length_7533_cov_3_4 | Penicillin/GL-7-ACA/AHL/aculeacin-A<br>acylase |
| 21 | 2.559 | ExploCarb_3808S_S2_1_length_789533_cov_14_559 | Peptidase S8, subtilisin-related |
| 22 | 2.527 | ExploCarb_3808S_S2_84_length_75002_cov_30_5 | PNPLA_dom |
| 25 | 2.390 | ExploCarb_3808S_S2_14352_length_1370_cov_1_2 | AB_hydrolase_1 |
| 42 | 1.597 | ExploCarb_3808S_S2_5211_length_3405_cov_171_3 | Peptidase family S66 |
| 44 | 1.491 | ExploCarb_3808S_S2_7708_length_2386_cov_3_1 | Lipase_bact_N |
| 58 | 1.269 | ExploCarb_3808S_S2_45_length_96604_cov_169_9 | PNPLA_dom |
| 69 | 1.142 | ExploCarb_3808S_S2_1180_length_13195_cov_3_10 | SGNH_hydro_sf |
| 84 | 1.047 | ExploCarb_3808S_S2_2714_length_6206_cov_211_1 | Penicillin/GL-7-ACA/AHL acylase |
| 89 | 1.016 | ExploCarb_3808S_S2_1272_length_12299_cov_60_1 | AB_hydrolase_1 |
| 104 | 0.911 | ExploCarb_3808S_S2_17847_length_1126_cov_1_2 | AB_hydrolase_1 |
| 116 | 0.819 | ExploCarb_3808S_S2_5424_length_3285_cov_7_1 | Peptidase S8, subtilisin-related |
| 139 | 0.710 | ExploCarb_3808S_S2_12631_length_1536_cov_2_2 | Peptidase S1C, Do |
| 167 | 0.583 | ExploCarb_3808S_S2_2577_length_6506_cov_183_2 | Peptidase S8, subtilisin-related |
| 189 | 0.521 | ExploCarb_3808S_S2_593_length_22580_cov_176_11 | Peptidase S8, subtilisin-related |
| 224 | 0.398 | ExploCarb_3808S_S2_4160_length_4192_cov_19_2 | Penicillin/GL-7-ACA/AHL acylase |
| 296 | 0.139 | ExploCarb_3808S_S2_21_length_132706_cov_167_47 | Peptidase S1C |
| 340 | 0.023 | ExploCarb_3808S_S2_858_length_17095_cov_20_14 | Hydrolase_4 (Serine aminopeptidase, S33) |

The active microbial community serine hydrolases enriched by eABPP comprised enzymes from all enzyme classes belonging to the serine hydrolase superfamily, including proteases (or peptidases as synonymous term), lipases, amidases and esterases^39^. Across both springs, we found proteins predicted as serine-type peptidases from the (super-)families S1 (chymotrypsin family, subfamily S1C), S8/S53 (clan SB: S8 (subtilisin family, including subfamily S8A), S53 (type peptidase: sedolisin)), S9 (prolyl oligopeptidase family), S15 (type peptidase: Xaa-Pro dipeptidyl peptidase), S33 (type peptidase: prolyl aminopeptidase), S45 (type peptidase: penicillin G acylase precursor), S49 (protease IV family), and S66 (type peptidase: murein tetrapeptidase LD-carboxypeptidase)^49^. Moreover, we detected proteins predicted as esterases/lipases, including enzymes from the SGNH hydrolase superfamily^50^ or enzymes containing a GDSL^51^, a PNPLA (patatin-like phospholipases)^52^ or a Lipase_bact_N^53^ domain, and a DUF915 family enzyme with structural similarity to esterases/lipases as well as a putative esterase from the UPF0227 family. Furthermore, enzymes predicted as alpha/beta fold-1 hydrolases^54^ without further classification but with esterases/lipases as structural homologs were identified from these datasets.

### Function validation of a selected serine hydrolase

We further confirmed the applicability of eABPP for the function-based protein identification from an environmental microbial community through subsequent bioinformatical and biochemical characterization of a selected serine hydrolase. From the list of predicted serine hydrolases identified by eABPP, we chose a putative esterase (ExploCarb_3811S_S4_483_length_13114_cov_941_5; Tab. 1) and retrieved and confined the corresponding gene sequence as described in the methods section. This enzyme was selected due to its relatively short and complete gene sequence as well as the availability of a straightforward enzyme kinetic assay.

The putative esterase is a 191 amino acid-containing protein possessing a UPF0227 domain (uncharacterized protein family 0227). A structural model of the putative esterase resembled secondary structural elements of characterized homologous esterases as well as its GTSLG sequence previously associated with thioesterase activity^55^ that fits within the conserved GxSxG motif of serine hydrolases (Figure 5a,b). The esterase sequence shows 100% identity to a predicted esterase (NCBI accession number: PMP76296.1) from a *Sulfurihydrogenibium* sp. MAG from a published Metagenome^48^, originating from the same spring. BLAST analysis predicted YqiA esterases, alpha/beta-fold hydrolases, other unspecified esterases and hypothetical proteins with no evidence of function at protein level as closely related hits. Structural analysis with HHpred reveals confident homology to a broad variety of esterases, including methylesterases, strigolactone esterases, thioesterases and carbohydrate esterases. Besides, structural homologs with diverging activities such as lipases, epoxide hydrolases, haloperoxidases, halogenalkane dehalogenases, polyethylene terephthalate hydrolases (PETases), cutinases, biphenyl catabolic (bph) hydrolases and other meta-cleavage product (MCP) hydrolases have been identified (Supplementary Tab. 3). Based on these bioinformatic analyses, the UPF0227 domain-containing serine hydrolase is most likely an esterase of the alpha/beta-hydrolases superfamily with unknown substrate specificity.

**Figure 5.**
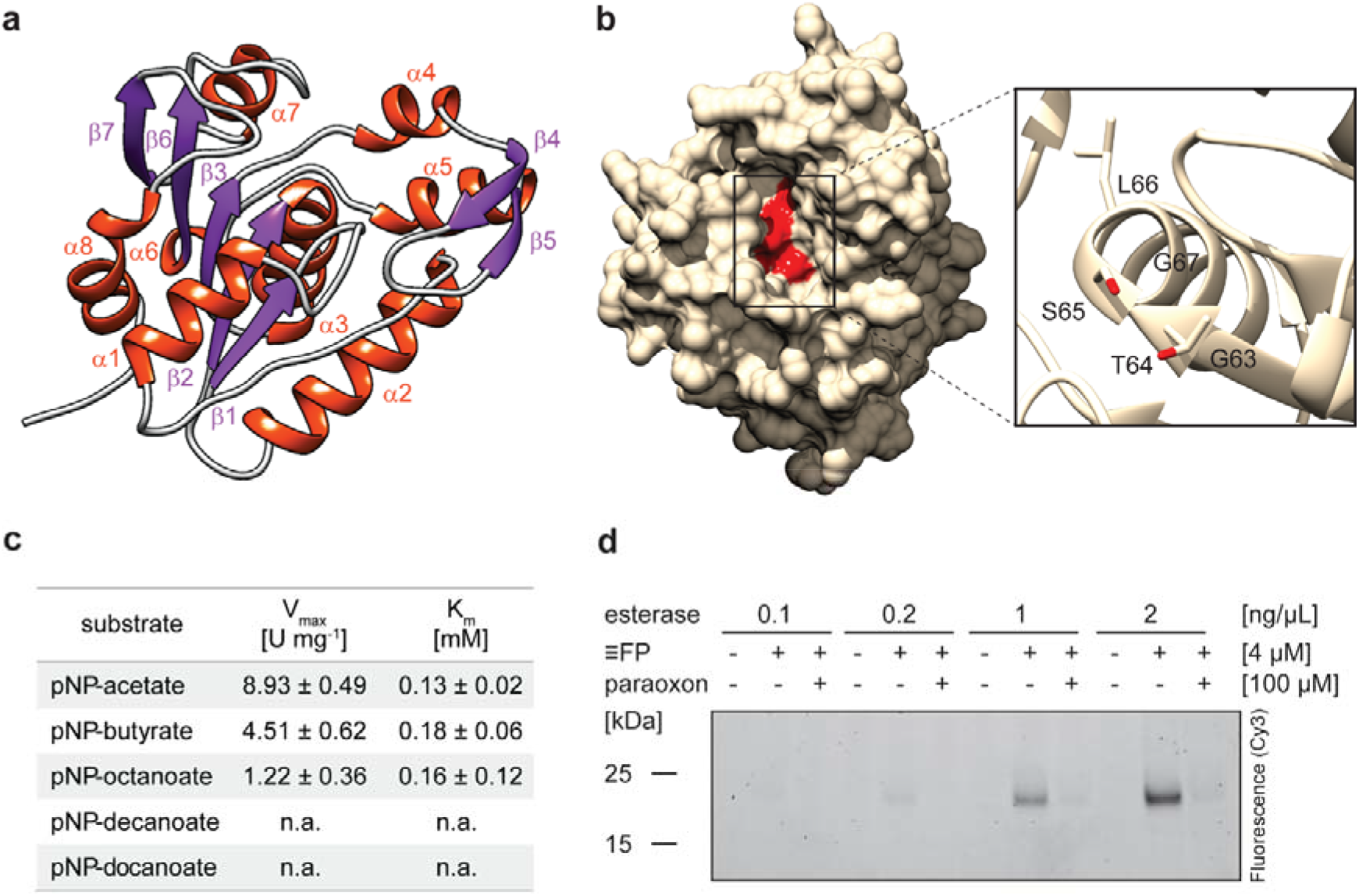
Characterization of the putative esterase. (**a**) Predicted structure of the UPF0227 protein with the secondary structures visualized in red (alpha helices) and purple (beta strands). The five-stranded parallel beta sheet consists of the strands β1, β2, β3, β6 and β7. (**b**) Surface-displaying structure of the putative esterase. The close-up displays the conserved serine hydrolase motif GxSxG, comprising the residues G63, T64, S65, L66 and G67, which is located in a substrate pocket (depicted in red). Structure prediction was performed with AlphaFold and the output was processed in Chimera. (**c**) Kinetic characterization of the esterase using the pNP-substrates pNP-acetate, -butyrate, -octanoate, -decanoate and -dodecanoate at concentrations up to 0.7 mM at pH 8.0 and 70°C and calculation of V_max_ and K_m_. Values represent the mean of three technical replicates ± SD. (**d**) *In vitro* labeling of varying amounts of the esterase with FP-alkyne in presence or absence of paraoxon.

In order to biochemically validate the function of the putative esterase, *E. coli* Rosetta (DE3) cells were transformed with the codon-optimized gene sequence cloned into a pET-28b(+) vector for heterologous protein expression. The purified enzyme was biochemically characterized using pNP-substrates. The putative esterase showed highest activity at a pH of 8.0 (Supplementary Fig. 2a) and a temperature of 70 °C (Supplementary Fig. 2b) using pNP-butyrate as a substrate. Moreover, enzyme kinetics were determined using different pNP-esters (Fig. 5c). Effective hydrolysis was observed for pNP-acetate (V_max_ = 8.93 U mg^−1^, K_m_ = 0.12 mM), pNP-butyrate (V_max_ = 4.51 U mg^−1^, K_m_ = 0.18 mM) and pNP-octanoate (V_max_ = 1.22 U mg^−1^, Km = 0.16 mM), confirming its function as an esterase, whereas no activity was measured for pNP-esters with longer chain lengths, such as pNP-decanoate or pNP-dodecanoate. Furthermore, *in vitro* ABPP of the purified esterase with FP-alkyne, analogously to the *in vivo* ABPP experiment, revealed robust labeling over a range of esterase concentrations, which could be strongly diminished by pre-incubation with the generic serine esterase inhibitor paraoxon (Fig. 5d). Consequently, the esterase was proven a bonafide target of the covalent-acting FP probe. In line with this result, the FP-alkyne was furthermore shown to reduce the esterase activity of the enzyme in correlation with the applied probe concentration (Supplementary Fig. 2c).

## Discussion

Extremal environments harbor microbial communities that provide myriad capabilities for identifying novel enzymatic activities and associated metabolic processes^56^. Due to their tremendous but mostly still hidden enzymatic potential, different strategies for their uncovering such as bioprospecting have emerged^11,12^. Despite recent technical advances, the study of the functional enzyme repertoire of an environmental microbial community and the reliable identification of the corresponding enzymes however remains a challenging task.

In this study, we established eABPP for the efficient function-based identification of active enzymes directly from the environment. The eABPP workflow relies on an activity-based labeling of a microbial community sample in combination with metagenome sequencing. In this way, the bioinformatic annotation of the metagenome can be directly confirmed through the activity-dependent reaction of the ABP-targeted enzyme, thereby providing direct experimental evidence for the bioinformatically predicted enzyme function along with its protein identification. So far, to the best of our knowledge, such a combination of metagenomics and MS-based ABPP has neither been established for protein identification from complex microbial communities and environmental samples, nor has it been applied directly in ecosystems.

As a showcase, we here reported the successful identification of active serine hydrolases from epi-sedimentary communities of two hot springs located in the Uzon Caldera region in Kamchatka (Russia), using the serine hydrolase target-specificity of FP-alkyne for functional enzyme annotation. Fluorophosphonate-based probes have been proven to show high specificity to serine hydrolases, even under harsh conditions such as high temperature^36,57^, providing information about the activity state of the labeled target enzymes across the examined microbial community. A robust LC-MS/MS workflow furthermore allows the identification and quantification of the respective proteins. Thorough bioinformatic analysis of the eABPP-enriched and -identified proteins from the sediments of the ‘Arkashin shurf’ (KAM3811) and the ‘Helicopter spring’ (KAM3808) revealed that most of the top hits (i.e. proteins with the highest log2-fold changes compared to control samples treated with DMSO) were indeed serine hydrolases, demonstrating the applicability of the method. It is therefore most likely that these proteins represent the most active serine hydrolases under the *in situ* conditions of the hot springs, due to either their higher abundances or their elevated activities compared to other serine hydrolases in the sampled environment. Of note, the applicability of our eABPP approach relies on different preconditions such as the availability of ABPs with sufficient target specificity and stability (e.g. in hot springs). It also required the development of distinct sample preparation procedures such as a suitable workflow for cleanup of proteins from the complex sample matrix prior to analysis by MS, or data analysis methodologies for detecting the low abundant single proteins from a metaproteome sample as well as the accurate construction of a metaproteome sequence database^29^.

To further corroborate the versatility of our eABPP approach, we heterologously expressed a putative esterase from the ‘uncharacterized protein family UPF0227’ which was found among the top 30 hits from KAM3811 enriched with FP-alkyne and demonstrated that this enzyme is efficiently hydrolyzing pNP-acetate (C2), pNP-butyrate (C4) and pNP-octanoate (C8), thereby confirming that the enzyme has esterase activity. Moreover, the esterase was efficiently labeled with FP-alkyne *in vitro*, thus supporting that the UPF0227 family protein indeed belongs to the serine hydrolases and overall confirming the eABPP-based identification (Fig. 5).

Overall, our study thus proved that the herein described workflow is suitable for the reliable identification of active serine hydrolases directly in the environment, by employing a well-studied ABP that has a broad spectrum of target enzymes. The proteins reported as hits for both springs include only sequences which could be readily identified as serine hydrolases, demonstrating the robustness of the applied approach. Please note that it is however likely that there are several more enzymes displaying serine hydrolase features and activities across the two sets of enriched proteins, which were not uncovered by this analysis.

The investigation of the functional potential of an environmental microbial community is, for example, used in the discovery of novel biocatalysts^12^. With the onset of the metagenomics era, the search for enzymes with prospect industrial use became far more effective^58^. Of note, most of the metagenomics-derived biocatalysts have been discovered by functional screening^14,59^. We thus believe that eABPP represents a novel approach for the efficient function-based identification of enzymes from microbial communities that may find application in industrial processes, thereby representing an alternative to functional metagenomics, which is quite time- and labor-intensive^15^, but frequently results in a rather low success rate^59^. Furthermore, our approach has the advantage that it allows to explore the biological diversity of active enzymes with dedicated functions from an ecosystem in their native state. With the majority of active serine hydrolases identified from our screen being annotated as subtilases (peptidase family S8) or penicillin acylases (peptidase family S15; Tab.1–2), we indeed detected thermostable representatives of the functionally highly diverse family of serine hydrolases that already find application for industrial purposes. In particular, proteases from the subtilisin family are utilized in the laundry industry^60^ or for the production of valuable chemicals from proteinaceous substrates^61,62^, whereas penicillin acylases are important enzymes for the production of 6-aminopenicillanic acid (6-APA), the β-lactam nucleus used in the manufacturing process of semi-synthetic antibiotics, through hydrolysis of natural penicillins^63^.

As a large and constantly growing repertoire of probes with a wide variety of warheads for targeting different enzymes or enzyme classes is already available, we believe that eABPP can be further extended to profile additional enzyme activities of microbial communities from diverse ecosystems, especially with respect to finding enzyme activities that harbor potential for application in industrial processes. These include, for example, cysteine proteases or glycoside hydrolases for which a broad set of probes is currently at hand^64,65^. Glycoside hydrolases, for instance, are of particular interest with regard to biocatalyst discovery since they largely function in (lignocellulosic) biomass degradation^66,67^. In addition, it might be reasonable to design more specific probes with activity towards serine hydrolases that function in the degradation of environmental pollutants (e. g. PETases, and different bph- and MCP hydrolases capable of cleaving mono- and bi-cyclic aromatic compounds) as these enzymes are of interest for environmental and industrial applications.

Finally, we believe that our approach may also find application besides industrial biocatalyst discovery. For example, we anticipate that eABPP may also be adapted for ecological research, for instance to decipher ecological enzyme activity patterns, e.g. from synergistic or mutual interactions, in microbial communities.

## Acknowledgements

BS, MK and IK acknowledge funding within the DFG-RSF Cooperation for joint German-Russian projects by the DFG (SI 642/12-1, KA 2894/6-1) and RSF (18-44-04024). This work was also supported by the DFG grant INST 20876/322-1 FUGG (to M. K. and F.K.).

## Author contributions

T.K. and T.V.K. performed in-field chemical labeling experiments, T.V.K. and I.V.K. performed DNA extraction and metagenomic sequencing, S.P.E. and A.J.P. performed refinement of metagenome data and metaproteome assembly, S.N., T.K. and L.S. performed protein extraction and affinity enrichment experiments, S.N. and F.K. performed MS analyses, T.K. performed protein expression and enzyme assays, B.S., M.K. and I.V.K. designed the workflow for the field ABPP, C.B, B.S. and M.K. supervised the study and S.N., T.K. B.S. and M.K. wrote the manuscript.

## References

1 Locey, K. J. & Lennon, J. T. Scaling laws predict global microbial diversity. Proc. Natl. Acad. Sci. USA 113, 5970–5975 (2016).

2 Verstraete, W. Microbial ecology and environmental biotechnology. ISME J. 1, 4–8 (2007).

3 Baquero, F., Coque, T. M., Galan, J. C. & Martinez, J. L. The Origin of Niches and Species in the Bacterial World. Front. Microbiol. 12, 657986 (2021).

4 Wohlgemuth, R. Biocatalysis - Key enabling tools from biocatalytic one-step and multi-step reactions to biocatalytic total synthesis. N. Biotechnol. 60, 113–123 (2021).

5 Elleuche, S., Schroder, C., Sahm, K. & Antranikian, G. Extremozymes - biocatalysts with unique properties from extremophilic microorganisms. Curr. Opin. Biotech. 29, 116–123 (2014).

6 Koskinen, P. E. P. et al. Bioprospecting thermophilic microorganisms from Icelandic hot springs for hydrogen and ethanol production. Energy Fuels 22, 134–140 (2008).

7 Sahoo, R. K., Kumar, M., Sukla, L. B. & Subudhi, E. Bioprospecting hot spring metagenome: lipase for the production of biodiesel. Environ. Sci. Pollut. Res. 24, 3802–3809 (2017).

8 Thankappan, S. et al. Bioprospecting thermophilic glycosyl hydrolases, from hot springs of Himachal Pradesh, for biomass valorization. AMB Expr. 8, 168 (2018).

9 Özdemir, S. C. & Uzel, A. Bioprospecting of hot springs and compost in West Anatolia regarding phytase producing thermophilic fungi. Sydowia 72, 1–11 (2020).

10 Prayogo, F. A. et al. Metagenomic applications in exploration and development of novel enzymes from nature: a review. J. Genet. Eng. Biotechnol. 18, 39 (2020).

11 Kennedy, J., Marchesi, J. R. & Dobson, A. D. W. Marine metagenomics: strategies for the discovery of novel enzymes with biotechnological applications from marine environments. Microb. Cell Fact. 7, 27 (2008).

12 DeCastro, M. E., Rodriguez-Belmonte, E. & Gonzalez-Siso, M. I. Metagenomics of Thermophiles with a Focus on Discovery of Novel Thermozymes. Front. Microbiol. 7, 1521 (2016).

13 Harrington, E. D. et al. Quantitative assessment of protein function prediction from metagenomics shotgun sequences. Proc. Natl. Acad. Sci. USA 104, 13913–13918 (2007).

14 Berini, F., Casciello, C., Marcone, G. L. & Marinelli, F. Metagenomics: novel enzymes from non-culturable microbes. FEMS Microbiol. Lett. 364, fnx211 (2017).

15 Lam, K. N., Cheng, J. J., Engel, K., Neufeld, J. D. & Charles, T. C. Current and future resources for functional metagenomics. Front. Microbiol. 6, 1196 (2015).

16 Cravatt, B. F., Wright, A. T. & Kozarich, J. W. Activity-based protein profiling: from enzyme chemistry to proteomic chemistry. Annu. Rev. Biochem. 77, 383–414 (2008).

17 Willems, L. I., Overkleeft, H. S. & van Kasteren, S. I. Current Developments in Activity-Based Protein Profiling. Bioconjugate Chem. 25, 1181–1191 (2014).

18 Yang, P. Y. & Liu, K. Activity-Based Protein Profiling: Recent Advances in Probe Development and Applications. Chembiochem 16, 712–724 (2015).

19 Fonović, M. & Bogyo, M. Activity-based probes as a tool for functional proteomic analysis of proteases. Expert Rev. Proteomics 5, 721–730 (2008).

20 Wright, M. H. & Sieber, S. A. Chemical proteomics approaches for identifying the cellular targets of natural products. Nat. Prod. Rep. 33, 681–708 (2016).

21 Barglow, K. T. & Cravatt, B. F. Activity-based protein profiling for the functional annotation of enzymes. Nat. Methods 4, 822–827 (2007).

22 Speers, A. E., Adam, G. C. & Cravatt, B. F. Activity-based protein profiling in vivo using a copper(I)-catalyzed azide-alkyne [3+2] cycloaddition. J. Am. Chem. Soc. 125, 4686–4687 (2003).

23 Speers, A. E. & Cravatt, B. F. Profiling enzyme activities in vivo using click chemistry methods. Chem. Biol. 11, 535–546 (2004).

24 Ovaa, H. et al. Chemistry in living cells: Detection of active proteasomes by a two-step labeling strategy. Angew. Chem. Int. Ed. 42, 3626–3629 (2003).

25 Berger, A. B., Vitorino, P. M. & Bogyo, M. Activity-based protein profiling: applications to biomarker discovery, in vivo imaging and drug discovery. Am. J. Pharmacogenomics 4, 371–381 (2004).

26 Keller, L. J., Babin, B. M., Lakemeyer, M. & Bogyo, M. Activity-based protein profiling in bacteria: Applications for identification of therapeutic targets and characterization of microbial communities. Curr. Opin. Chem. Biol. 54, 45–53 (2020).

27 Morimoto, K. & van der Hoorn, R. A. The Increasing Impact of Activity-Based Protein Profiling in Plant Science. Plant Cell Physiol. 57, 446–461 (2016).

28 Klaus, T. et al. Activity-Based Protein Profiling for the Identification of Novel Carbohydrate-Active Enzymes Involved in Xylan Degradation in the Hyperthermophilic Euryarchaeon Thermococcus sp. Strain 2319×1E. Front. Microbiol. 12, 734039 (2021).

29 Whidbey, C. & Wright, A. T. Activity-Based Protein Profiling-Enabling Multimodal Functional Studies of Microbial Communities. Curr. Top. Microbiol. Immunol. 420, 1–21 (2019).

30 Mayers, M. D., Moon, C., Stupp, G. S., Su, A. I. & Wolan, D. W. Quantitative Metaproteomics and Activity-Based Probe Enrichment Reveals Significant Alterations in Protein Expression from a Mouse Model of Inflammatory Bowel Disease. J. Proteome Res. 16, 1014–1026 (2017).

31 Chatterjee, S. et al. A comprehensive and scalable database search system for metaproteomics. BMC Genomics 17, 642 (2016).

32 Chauvigne-Hines, L. M. et al. Suite of activity-based probes for cellulose-degrading enzymes. J. Am. Chem. Soc. 134, 20521–20532 (2012).

33 Ansong, C. et al. Characterization of protein redox dynamics induced during light-to-dark transitions and nutrient limitation in cyanobacteria. Front. Microbiol. 5, 325 (2014).

34 Sadler, N. C. et al. Live cell chemical profiling of temporal redox dynamics in a photoautotrophic cyanobacterium. ACS Chem. Biol. 9, 291–300 (2014).

35 Bennett, K., Sadler, N. C., Wright, A. T., Yeager, C. & Hyman, M. R. Activity-Based Protein Profiling of Ammonia Monooxygenase in Nitrosomonas europaea. Appl. Environ. Microbiol. 82, 2270–2279 (2016).

36 Zweerink, S. et al. Activity-based protein profiling as a robust method for enzyme identification and screening in extremophilic Archaea. Nat. Commun. 8 (2017).

37 Sakoula, D. et al. Universal activity-based labeling method for ammonia- and alkane-oxidizing bacteria. ISME J. 16, 958–971 (2022).

38 Liu, Y. S., Patricelli, M. P. & Cravatt, B. F. Activity-based protein profiling: The serine hydrolases. Proc. Natl. Acad. Sci. USA 96, 14694–14699 (1999).

39 Simon, G. M. & Cravatt, B. F. Activity-based Proteomics of Enzyme Superfamilies: Serine Hydrolases as a Case Study. J. Biol. Chem. 285, 11051–11055 (2010).

40 Chandra, P., Enespa, Singh, R. & Arora, P. K. Microbial lipases and their industrial applications: a comprehensive review. Microb. Cell Fact. 19, 169 (2020).

41 Panda, T. & Gowrishankar, B. S. Production and applications of esterases. Appl. Microbiol. Biotechnol. 67, 160–169 (2005).

42 Barzkar, N. et al. Marine Bacterial Esterases: Emerging Biocatalysts for Industrial Applications. Appl. Biochem. Biotechnol. 193, 1187–1214 (2021).

43 Anobom, C. D. et al. From Structure to Catalysis: Recent Developments in the Biotechnological Applications of Lipases. Biomed. Res. Int. 2014, 684506 (2014).

44 Romano, D. et al. Esterases as stereoselective biocatalysts. Biotechnol. Adv. 33, 547–565 (2015).

45 Wu, Z. M., Liu, C. F., Zhang, Z. Y., Zheng, R. C. & Zheng, Y. G. Amidase as a versatile tool in amide-bond cleavage: From molecular features to biotechnological applications. Biotechnol. Adv. 43 (2020).

46 Pentecost, A., Jones, B. & Renaut, R. W. What is a hot spring? Can. J. Earth Sci. 40, 1443–1446 (2003).

47 Burgess, E. A., Unrine, J. M., Mills, G. L., Romanek, C. S. & Wiegel, J. Comparative geochemical and microbiological characterization of two thermal pools in the Uzon Caldera, Kamchatka, Russia. Microb. Ecol. 63, 471–489 (2012).

48 Wilkins, L. G. E., Ettinger, C. L., Jospin, G. & Eisen, J. A. Metagenome-assembled genomes provide new insight into the microbial diversity of two thermal pools in Kamchatka, Russia. Sci. Rep. 9, 3059 (2019).

49 Rawlings, N. D. et al. The MEROPS database of proteolytic enzymes, their substrates and inhibitors in 2017 and a comparison with peptidases in the PANTHER database. Nucleic Acids Res. 46, D624–D632 (2018).

50 Mølgaard, A., Kauppinen, S. & Larsen, S. Rhamnogalacturonan acetylesterase elucidates the structure and function of a new family of hydrolases. Structure 8, 373–383 (2000).

51 Akoh, C. C., Lee, G. C., Liaw, Y. C., Huang, T. H. & Shaw, J. F. GDSL family of serine esterases/lipases. Prog. Lipid Res. 43, 534–552 (2004).

52 Baulande, S. & Langlois, C. [Proteins sharing PNPLA domain, a new family of enzymes regulating lipid metabolism]. Med. Sci. 26, 177–184 (2010).

53 Chuang, Y. C., Chiou, S. F., Su, J. H., Wu, M. L. & Chang, M. C. Molecular analysis and expression of the extracellular lipase of Aeromonas hydrophila MCC-2. Microbiol. 143, 803–812 (1997).

54 Ollis, D. L. et al. The Alpha/Beta-Hydrolase Fold. Protein Eng. 5, 197–211 (1992).

55 Thankachan, D. et al. A trans-Acting Cyclase Offloading Strategy for Nonribosomal Peptide Synthetases. ACS Chem. Biol. 14, 845–849 (2019).

56 Shu, W. S. & Huang, L. N. Microbial diversity in extreme environments. Nat. Rev. Microbiol. 20, 219–235 (2022).

57 Kidd, D., Liu, Y. & Cravatt, B. F. Profiling serine hydrolase activities in complex proteomes. Biochemistry 40, 4005–4015 (2001).

58 Ferrer, M. et al. Estimating the success of enzyme bioprospecting through metagenomics: current status and future trends. Microb. Biotechnol. 9, 22–34 (2016).

59 Zhang, L. et al. Advances in Metagenomics and Its Application in Environmental Microorganisms. Front. Microbiol. 12, 766364 (2021).

60 Niehaus, F. et al. Enzymes for the laundry industries: tapping the vast metagenomic pool of alkaline proteases. Microb. Biotechnol. 4, 767–776 (2011).

61 De Oliveira Martinez, J. P. et al. Challenges and Opportunities in Identifying and Characterising Keratinases for Value-Added Peptide Production. Catalysts 10, 184 (2020).

62 Li, J. et al. A Novel Gelatinase from Marine Flocculibacter collagenilyticus SM1988: Characterization and Potential Application in Collagen Oligopeptide-Rich Hydrolysate Preparation. Mar. Drugs 20, 48 (2022).

63 Arroyo, M., de la Mata, I., Acebal, C. & Castillon, M. P. Biotechnological applications of penicillin acylases: state-of-the-art. Appl. Microbiol. Biotechnol. 60, 507–514 (2003).

64 Wu, L. et al. An overview of activity-based probes for glycosidases. Curr. Opin. Chem. Biol. 53, 25–36 (2019).

65 Serim, S., Haedke, U. & Verhelst, S. H. Activity-based probes for the study of proteases: recent advances and developments. ChemMedChem 7, 1146–1159 (2012).

66 Suleiman, M., Kruger, A. & Antranikian, G. Biomass-degrading glycoside hydrolases of archaeal origin. Biotechnol. Biofuels 13 (2020).

67 Prieto, A. et al. Fungal glycosyl hydrolases for sustainable plant biomass valorization: Talaromyces amestolkiae as a model fungus. Int. Microbiol. 24, 545–558 (2021).

