## Supplementary Information for "Environmental activity-based protein profiling for function-driven enzyme discovery from natural communities"

### Supplementary Figures

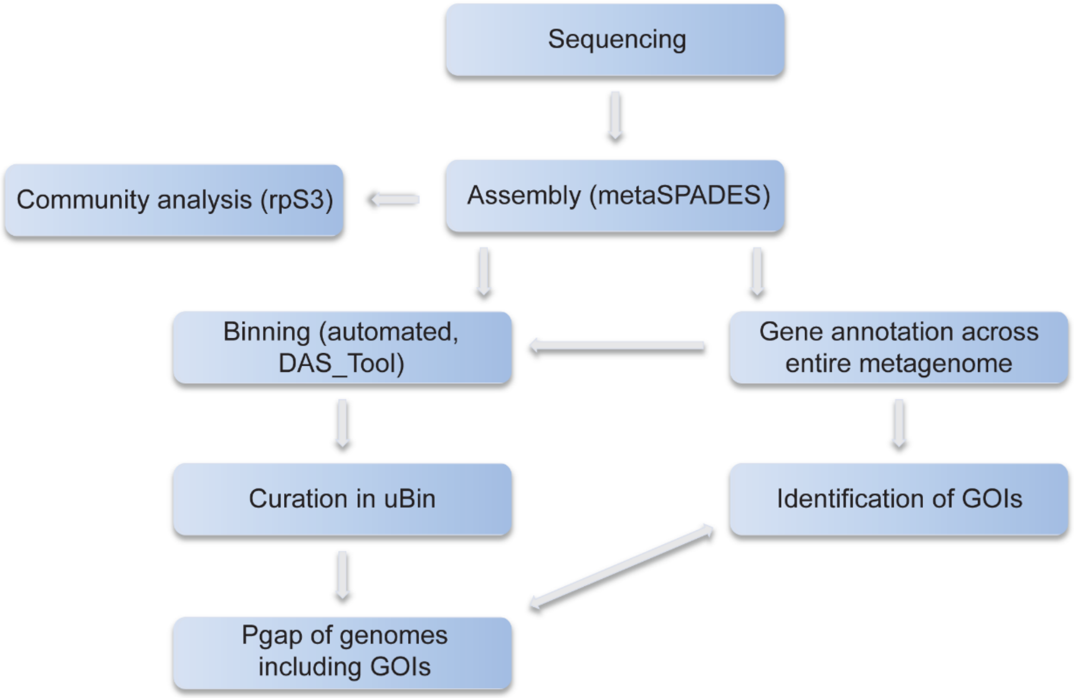


**Supplementary Figure 1. Bioinformatics pipeline**

Visualization of the bioinformatics pipeline described in the Method section. The pipeline starts with sequencing, followed by quality-filtering and assembly, to building the metaproteome database and the identification of genes of interest.

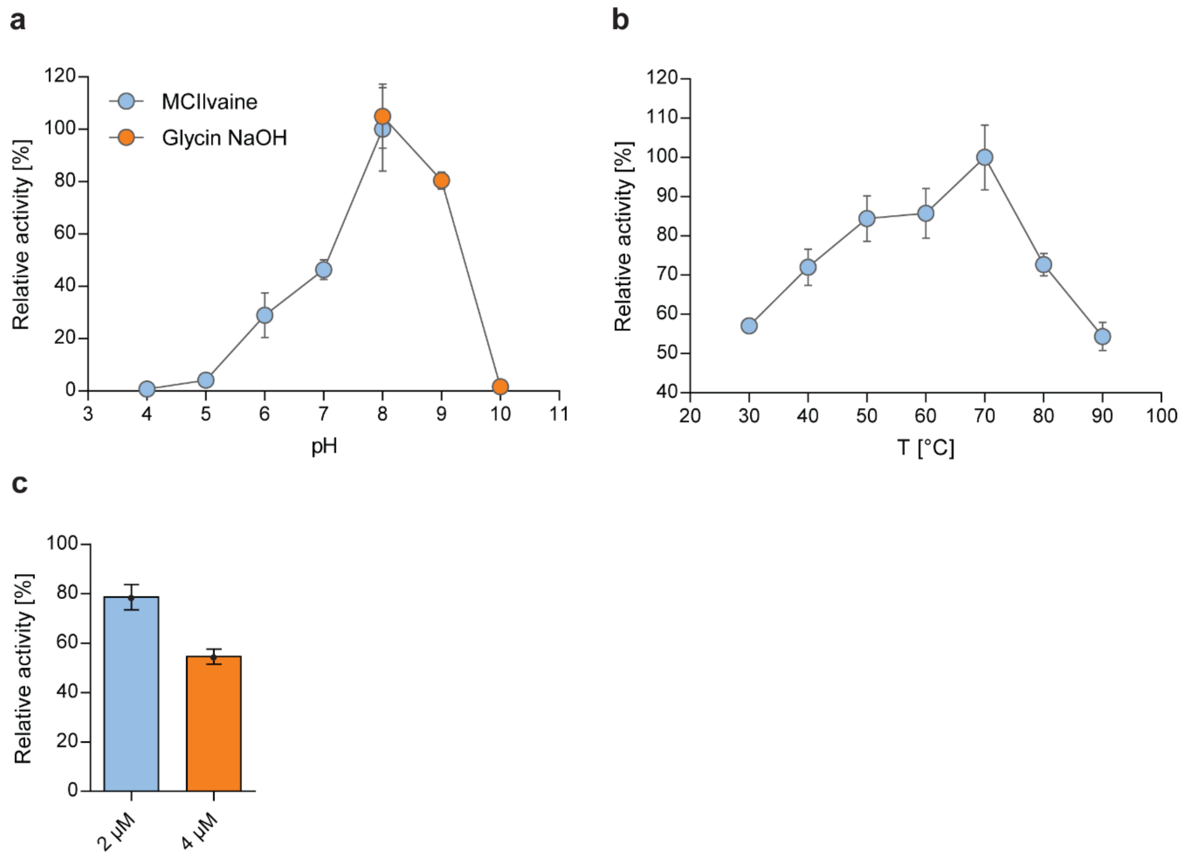


**Supplementary Figure 2. Biochemical characterization of the heterologously expressed esterase**

Effect of pH (**a**) and temperature (**b**) on the relative activity of the esterase with pNP-butyrate. (**c**) Activity inhibition of the esterase upon preincubation with 2 µM or 4 µM FP-alkyne at pH 7.5 and 70° C for 10 min. The remaining activity was measured using pNP-acetate as the substrate. Error bars represent SD of three technical replicates.

### Supplementary Tables

**Supplementary Table 1.** Relative abundance of microorganisms in KAM3811 based on the coverage of scaffolds containing the respective rpS3 gene.

| Nearest predicted relative based on BLASTp | Phylum | Domain | Abundance (rpS3) |
|---|---|---|---|
| Sulfurihydrogenibium sp. | Aquificota | Bacteria | 2329.1 |
| Pyrobaculum ferrireducens | Thermoproteota | Archaea | 411.3 |
| Caldisphaera sp. | Thermoproteota | Archaea | 389.6 |
| Hydrogenobaculum sp. | Aquificota | Bacteria | 281.7 |
| Nitrososphaera sp. | Thermoproteota | Archaea | 160.8 |
| Sulfolobales (SCGC AB-777 J03) | Thermoproteota | Archaea | 158.5 |
| Aciduliprofundum sp. | Thermoplasmatota | Archaea | 139.0 |
| Caldisericum exile | Caldisericota | Bacteria | 78.2 |
| Fervidicoccus sp. | Thermoproteota | Archaea | 59.7 |
| Caldimicrobium thiodismutans | Desulfobacterota | Bacteria | 58.9 |
| Candidatus Nanobsidianus stetteri | Nanoarchaeota | Archaea | 38.5 |
| Thermoproteus sp. (JCHS 4) | Thermoproteota | Archaea | 26.9 |
| Aciduliprofundum sp. | Thermoplasmatota | Archaea | 19.1 |
| Caldivirga maquilingensis (IC-167) | Thermoproteota | Archaea | 17.1 |
| Mesoaciditoga sp. | Thermotogota | Bacteria | 13.9 |
| Caldivirga maquilingensis (IC-167) | Thermoproteota | Archaea | 11.5 |
| Thiomonas sp. (CB3) | Proteobacteria | Bacteria | 11.1 |
| Thiomonas sp. (20-64-5) | Proteobacteria | Bacteria | 10.9 |
| Acidilobus saccharovorans (345-15) | Thermoproteota | Archaea | 7.7 |

**Supplementary Table 2.** Relative abundance of microorganisms in KAM3808 based on the coverage of the scaffold containing the respective rpS3 gene.

| **Nearest predicted relative based on BLASTp** | **Phylum** | **Domain** | **Abundance (rpS3)** |
|---|---|---|---|
| Aciduliprofundum sp. | Thermoplasmatota | Archaea | 488.1 |
| Caldisericum exile | Caldisericota | Bacteria | 408.2 |
| Caldimicrobium thiodismutans | Desulfobacterota | Bacteria | 139.9 |
| Archaeon, unclassified | not predicted | Archaea | 124.6 |
| Sulfurihydrogenibium sp. | Aquificota | Bacteria | 119.9 |
| Nitrososphaera sp. | Thermoproteota | Archaea | 93.7 |
| Thermus arciformis | Deinococcota | Bacteria | 81.6 |
| Candidate division WOR-3 JGI Cruoil (03_51_56) | WOR-3 | Bacteria | 64.5 |
| Dictyoglomus turgidum (DSM 6724) | Dictyoglomota | Bacteria | 54.9 |
| Candidatus Micrarchaeota (CG1_02_51_15) | Micrarchaeota | Archaea | 52.9 |
| Thermoflexia | Chloroflexota | Bacteria | 46.3 |
| Chlorobi (MS-B_bin-24) | Bacteroidota | Bacteria | 37.1 |
| Parcubacteria (ADurb.Bin305) | Patescibacteria | Bacteria | 31.5 |
| Candidatus Bathyarchaeota (ex4484_205) | Thermoproteota | Archaea | 25.4 |
| Aciduliprofundum sp. | Thermoplasmatota | Archaea | 20.1 |
| Thermosulfidibacter takaii (ABI70S6) | Thermosulfidibacterota | Bacteria | 18.7 |
| Fervidicoccus sp. | Thermoproteota | Archaea | 18.1 |
| Thermodesulfovibrio aggregans | Nitrospirota | Bacteria | 12.8 |
| Bacterium, unclassified | not predicted | Bacteria | 11.7 |
| Candidatus Cryosericum odellii | Caldisericota | Bacteria | 11.7 |
| Archaeon, unclassified | not predicted | Archaea | 11.6 |
| Thermogutta terrifontis | Planctomycetota | Bacteria | 10.7 |
| Carboxydocella sp. | Firmicutes | Bacteria | 10.5 |
| Candidatus Woesearchaeota (ex4484_78) | Nanoarchaeota | Archaea | 9.7 |
| Bacterium, unclassified | not predicted | Bacteria | 9.5 |
| Candidatus Roizmanbacteria | Patescibacteria | Bacteria | 9.1 |
| Elusimicrobia (CG08_land_8_20_14_0_20_51_18) | Elusimicrobiota | Bacteria | 8.1 |
| Archaeon, unclassified | not predicted | Archaea | 7.8 |
| Verrucomicrobia | Verrucomicrobiota | Bacteria | 7.8 |
| Candidate division Zixibacteria (RBG_16_43_9) | Zixibacteria | Bacteria | 7.6 |
| Candidatus Omnitrophica | Omnitrophota | Bacteria | 7.4 |
| Candidatus Aenigmarchaeota | Aenigmatarchaeota | Archaea | 7.2 |
| Geobacter sp. | Desulfobacterota | Bacteria | 6.8 |
| Bacterium HR16 | Armatimonadota | Bacteria | 6.7 |
| Heliorestis acidaminivorans | Firmicutes | Bacteria | 6.5 |
| Archaeon, unclassified | not predicted | Archaea | 6 |
| Thermofilum uzonense | Thermoproteota | Archaea | 5.8 |
| Fervidobacterium nodosum (Rt17-B1) | Thermotogota | Bacteria | 5.4 |
| Archaeon, unclassified | not predicted | Archaea | 5.3 |
| Thermodesulforhabdus norvegica | Desulfobacterota | Bacteria | 5.2 |
| Archaeon, unclassified | not predicted | Archaea | 5.1 |
| Candidatus Atribacteria (ADurb.Bin276) | Atribacterota | Bacteria | 5.1 |
| Desulfurella sp. | Campylobacterota | Bacteria | 5.1 |
| Ignavibacteriales (CG07_land_8_20_14_0_80_59_12) | Bacteroidota | Bacteria | 4.3 |

| | | | |
|---|---|---|---|
| Parcubacteria (ADurb.Bin305) | Patescibacteria | Bacteria | 4 |
| Candidatus Aminicenantes | Acidobacteriota | Bacteria | 3.4 |
| Candidatus Aminicenantes | Acidobacteriota | Bacteria | 3.4 |
| Lentisphaeria | Verrucomicrobiota | Bacteria | 3.4 |
| Ignavibacteria (RBG_13_36_8) | Bacteroidota | Bacteria | 2.7 |

**Supplementary Table 3.** Selected structural homologs of the putative esterase determined with HHpred.

| Hit | Function | Origin | Probability [%] | E-value | ScoreOrigin |
|---|---|---|---|---|---|
| 4FLE_A | Ancestral haloalkane dehalogenase AncHLD3; | Synthetic construct | 99.96 | 7.8e-26 | 140.97 |
| 413F_A | serine hydrolase CCSP0084; MCP cleaving | *Cycloplasticus* sp. P1 | 99.93 | 4.3e-24 | 141.6 |
| 3N98_A | Chloroperoxidase F; Haloperoxidase, Oxidoreductase | *Pseudomonas fluorescens* | 99.93 | 3.5e-24 | 138.8 |
| 3G9X_A | Haloalkane dehalogenase; alpha/beta hydrolase, helical cap domain | *Rhodococcus rhodochrous* | 99.93 | 1.6e-23 | 137.84 |
| 4UHC_A | Esterase; alpha/beta hydrolase, pnp-ester cleaving | *Thermogutta terrifontis* | 99.93 | 5.4e-24 | 138.98 |
| 3RM3_A | Thermostable monoacylglycerol lipase; alpha/beta hydrolase fold | *Bacillus* sp. H257 | 99.92 | 1.5e-23 | 136.84 |
| 3PFB_B | Cinnamoyl esterase; alpha/beta hydrolase fold, esterase, hydrolase, cinnamoyl/feruloyl esterase | *Lactobacillus johnsonii* | 99.92 | 7.7e-23 | 133.82 |
| 1C4X_A | 2-hydroy-6-oxo-6-phenylhexa-2,4-dienoate hydrolase (BPHD); PCB degradation | *Rhodococcus* sp. Strain Rha1 | 99.92 | 7.9e-24 | 138.27 |
| 5OLU_A | carboxyl esterase, 1,2-O-isopropylideneglycerol hydrolyzing, lipase, alpha/beta hydrolase | *Bacillus coagulans* | 99.92 | 1.3e-22 | 136.85 |
| 4LXH_A | MCP Hydrolase; carbon-carbon bond hydrolase, Rossmann Fold, alpha/beta hydrolase fold | *Sphingomonas wittichii* RW1 | 99.92 | 2.2e-23 | 135.76 |
| 5Y6Y_B | Epoxide hydrolase | *Vigna radiata* | 99.92 | 2.1e-23 | 138.9 |
| 2WTM_C | Promiscuous Feruloyl Esterase (Est1E) | *Butyrivibrio Proteoclasticus* | 99.92 | 1.2e-22 | 130.56 |
| 4C6H_A | Haloalkane dehalogenase | Rhodobacteraceae | 99.92 | 5.1e-23 | 135.33 |
| 2XTO_A | Haloalkane Dehalogenase | *Plesiocystis pacifica* SIR-I | 99.92 | 3.5e-23 | 136.21 |
| 1Q0R_A | aclacinomycin methylesterase; Anthracycline, methylesterase, polyketide hydrolase | *Serratia marcescens* | 99.92 | 9.6e-24 | 139.12 |
| 6F9O_A | haloalkane dehalogenase DpcA | *Psychrobacter cryohalolentis* K5 | 99.92 | 5.3e-23 | 136.58 |

| | | | | | |
|---|---|---|---|---|---|
| 6Y9G_B | Ancestral haloalkane dehalogenase AncHLD5 | Synthetic construct | 99.91 | 1.5e-22 | 133.85 |
| 2E3J_A | epoxide hydrolase B (Rv1938) | *mycobacterium tuberculosis* | 99.91 | 1.3e-22 | 137.43 |
| 5XKS_F | Thermostable monoacylglycerol lipase | *Geobacillus* sp. 12AMOR | 99.91 | 4.6e-22 | 128.73 |
| 2WFL_A | polyneuridine aldehyde esterase (PNAE) | *Rauvolfia serpentina* | 99.91 | 3.3e-22 | 128.71 |
| 3BF7_B | Esterase YbfF; esterase, thioesterase | *Escherichia coli* | 99.91 | 2.4e-22 | 128.91 |
| 6THS_A | Serine esterase, cutinase S165A | Uncultured bacterium | 99.9 | 6.5e-22 | 129 |
| 5XWZ_B | Alpha/beta-hydrolase, lactonase, zearalenone hydrolase | *Cladophialophora bantiana* | 99.9 | 3.8e-22 | 130.52 |
| 1IUP_A | meta-Cleavage product hydrolase; aromatic compounds, cumene, isopropylbenzene, meta-cleavage compound hydrolase | *Pseudomonas fluorescens* IP01 (CumD) | 99.9 | 1.6e-21 | 127.29 |
| 6BA9_A | yersiniabactin synthesis enzyme, YbtT; Thioesterase, non-ribosomal peptide synthesis | *Escherichia coli* | 99.9 | 3e-22 | 129.51 |
| 4CCY_A | Carboxylesterase YBFK, naproxenmethylester hydrolase | *Bacillus subtilis* | 99.9 | 6.4e-23 | 135.19 |
| 2WUE_B | 2-hydroy-6-oxo-6-phenylhexa-2,4-dienoate hydrolase (BPHD | *Mycobacterium tuberculosis* | 99.89 | 2e-21 | 127.37 |
| 2RHW_A | BphD, C-C Bond Hydrolase Involved in Polychlorinated Biphenyls Degradation | *Burkholderia xenovorans* LB400 | 99.89 | 1.1e-21 | 127.92 |
| 3FCY_A | Xylan esterase 1; alpha/beta hydrolase, carbohydrate esterase, CE7 | *Thermoanaerobacterium* sp. JW/SL YS485 | 99.89 | 8.9e-22 | 133.37 |
| 6AGQ_A | acetyl xylan esterase | *Paenibacillus* sp. R4 | 99.89 | 1.8e-21 | 130.01 |
| 1MJ5_A | 1,3,4,6-tetrachloro-1,4-cyclohexadiene hydrolase; LINB, hydrolase, Haloalkane dehalogenase | *Sphingomonas paucimobilis* UT26 | 99.88 | 1e-20 | 125.53 |
| 5XH2_A | Poly(ethylene terephthalate) hydrolase | *Ideonella sakaiensis* 201-F6 | 99.88 | 5.6e-21 | 124.45 |
| 4CG1_A | Cutinase, PET degrading hydrolase | *Thermobifida fusca* | 99.86 | 1.4e-19 | 119.52 |
| 7NEI_B | Polyester Hydrolase Leipzig 7 (PHL-7); PETase, Cutinase | unidentified | 99.86 | 7.4e-20 | 119.78 |

### Material and Methods

#### Sample collection and chemical labeling

Sediments for chemical labeling experiments have been sampled from the two hot springs 'Arkashin shurf' (54°30.0016N 160°00.2021E, 65.5-72° C, pH = 5.01, internal number #3811) and 'Helicopter spring' (54°30.003N 160°00.4375E, 58.1° C, pH = 5.62, internal number #3808), both located at the Uzon volcanic caldera (Kamchatka Peninsula, Russia). FP-alkyne was dissolved in DMSO. A slurry of sediments in spring water was collected and after gentle mixing, 10 ml of the slurry was dispensed in a reaction tube and incubated with 4 µM FP-alkyne for 2 h with occasional shaking while being placed back in the spring. An equal volume of DMSO was added to the negative controls. All samples were prepared in triplicates. For metagenomic sequencing, an unlabeled aliquot of the slurry was prepared. For downstream processing, the samples were transported to the laboratory on dry ice, the slurry was centrifuged to remove the spring water (12,000 × *g*, room temperature, 10 min) and the sediments were stored at -20° C until further processing.

#### DNA extraction and metagenomic sequencing

Total DNA was isolated from the sediments using phenol-chloroform extraction as described in Gavrilov *et al.*[1]. Prior to isolation, the cells were disrupted using a series of freezing-thawing cycles. Concentration of DNA was measured on a Qubit 2.0 fluorometer (Invitrogen, USA). Shotgun metagenome library preparation and sequencing were done at BioSpark Ltd., Moscow, Russia. The KAPA HyperPlus Library Preparation Kit (KAPA Biosystems, USA) was used for library construction according to the manufacturer's protocol and sequencing was perfromed on a NovaSeq 6000 platform (Illumina, USA) with the NovaSeq 6000 S2 Reagent Kit, which can read 100 nucleotides from each end (200 cycles).

#### Genome-resolved metagenomics

Raw reads were quality filtered and cleaned with BBDuk (https://jgi.doe.gov/data-and-tools/bbtools) and sickle[2] and followed by assembly with metaSPAdes[3] (version 3.14). Assembled sequences below 1 kbp in length were discarded and gene prediction was carried out using Prodigal in meta mode[4] followed by annotation against the FunTaxDB[5], which is based on the UniRef100[6] database. Community composition of samples was determined based on ribosomal protein S3 (rpS3) and its respective coverage on scaffolds in the individual metagenomes. The coverage was determine via mapping[7] of metagenomic reads and taken as the relative abundance of rpS3 genes and their respective microbes in the community. Binning of genomes was performed with MaxBin2[8], ABAWACA[9] and emergent self-organizing maps[10]. High quality genomes were identified using DAS Tool[11] and further curated with uBin[5]. A phylogenetic tree was calculated with GTDB-Tk[12] based on the dereplicated genomes. Genome completeness, contamination, GC content and genome length was calculated via CheckM[13]. The target genome that contained the serine hydrolase chosen for biochemical characterization (see Heterologous protein expression and purification) was analyzed and re-annotated with NCBI's PGAP[14] (Prokaryotic Genomes Annotation Pipeline) to improve start and end of gene prediction.

**Protein extraction and clean-up**

For protein extraction from labeled sediments, the collected organic matter was thawed on ice and taken up in 5 mL of extraction buffer (100 mM Tris-HCl pH 8.8, 0.1 M DTT, 50 mM EDTA, 1.5% SDS, 30% sucrose). The cells were lysed in a three-step sonication procedure: seven iterations of sonication in an ultrasonic bath (BANDELIN electronic, Germany) for 1 min followed by vigorous mixing, 10 cycles with high power in a Bioruptor UCD-200 (Diagenode, Belgium) device with the following conditions: 1 min pulse and 30 sec pause and another ten iterations of sonication in an ultrasonic bath as described before. The extracts were then cleared by centrifugation (100 × *g*, room temperature, 5 min) and the supernatant was collected in a fresh tube. The debris were centrifuged again (15,000 × *g*, room temperature, 20 min) and the supernatant was combined with the supernatant in the fresh tube. The combined supernatant was subjected to another centrifugation step (15,000 × *g*, room temperature, 15 min) to separate the soluble protein containing fraction from any remaining debris. Protein clean-up was done by performing a phenol extraction[15] with downstream ammonium acetate precipitation[16] according to the literature with few modifications. In brief, the protein containing fractions were mixed and incubated (15 min, room temperature, shaking) with an equal volume of TE-buffered liquid Phenol (Carl Roth, Germany; #0038). To achieve phase separation, the samples were centrifuged (12,000 × *g*, room temperature, 10 min). The upper phenol phase was collected and re-extracted with extraction buffer thrice. Thereto, the phenol phase was mixed with an equal volume of extraction buffer and the phases were separated by centrifugation (12,000 × *g*, room temperature, 10 min). The proteins in the phenol phase were precipitated with a five-fold volume of 0.1 M ammonium acetate in methanol (-20 °C, overnight). The proteins were collected by centrifugation (12,000 × *g*, 4 °C, 30 min) and the pellet was successively washed twice with 0.1 M ammonium acetate in methanol, twice with 80% (v/v) acetone, and once with 70% (v/v) ethanol, respectively. Each washing step included an incubation step at -20 °C for 20-30 min prior to centrifugation (12,000 × *g*, 4 °C, 30 min). The precipitated proteins were dissolved in 100 µL 8 M urea in 50 mM $HNa_2PO_4$, pH 8.0 and further diluted with 50 mM $HNa_2PO_4$ pH 8.0 to a final concentration of 2 M urea. The protein concentration of the resulting protein solutions was determined by a modified Bradford assay with Roti-Nanoquant (Carl Roth, Germany).

**Click reaction and affinity purification**

400-650 µg of total protein were subjected to a click reaction with 10 µM 5/6-TAMRA-biotin-$N_3$ (Jena Bioscience, Germany; #CLK-1048), 100 µM TBTA, 2 mM TCEP and 2 mM $CuSO_4$ (all purchased from Sigma-Aldrich, USA) in a total reaction volume of 500 µL (1 h, room temperature, in the dark). Prior to affinity purification, unbound reporter and salts from the click reaction were removed by methanol-chloroform precipitation[17]. The resulting protein pellet was air-dried and subsequently dissolved (37 °C, ~1 h) in 850 µL 2% (w/v) SDS in 1× PBS (155 mM NaCl, 3 mM $Na_2HPO_4$, 1.06 mM $KH_2PO_4$, pH 7.4). Insoluble particles were removed by centrifugation (21,000 × *g*, 37 °C, 5 min) and the cleared protein solution was diluted with 1× PBS to a final concentration of 0.2% (w/v) SDS. The obtained protein mixture was incubated with 100 µl of pre-equilibrated avidin beads slurry (Thermo Scientific, USA; #20219) while gently tumbling (~1h, room temperature, in the dark). Subsequently, the beads were washed five times with 10 mL 1% (w/v) SDS (10 min, room temperature, gently rotating) and collected by centrifugation (400 × *g*, 5 min). To remove SDS from the samples, the beads were then washed four times with 1 mL of ultrapure water (VWR Chemicals, USA; 5 min, room temperature, vigorously shaking) and collected by centrifugation (3,000 × *g*, 1 min).

**On-bead digestion of captured proteins**

After affinity enrichment, the beads were taken up in 100 µL 0.8 M urea in 50 mM ammonium bicarbonate (ABC). Disulfide bonds were reduced by adding 10 mM DTT (Sigma-Aldrich, USA) in 50 mM ABC (1 h, room temperature, vigorous shaking) and the generated cysteine mercapto groups were masked by alkylation with 25 mM Iodoacetamide (IAM; Sigma-Aldrich, USA) in 50 mM ABC (1 h, room temperature, in the dark, vigorous shaking). Excess IAM was then quenched by adding DTT (final concentration of 35 mM, 10 min, room temperature, vigorous shaking). Protein on-bead digestion was started by adding 1 µg Trypsin (Thermo Scientific, USA; #90057) dissolved in 50 mM acetic acid (37 °C, ~16 h, vigorous shaking). After digestion, the reaction solution was cleared by centrifugation (3,000 × *g*, room temperature, 5 min) and the supernatant (contains the digestion products (peptides)) was transferred to a fresh reaction vessel and the digestion reaction stopped by adding formic acid (FA) to a final concentration of 5% (v/v). Next, the beads were washed with 50 µL 1% (v/v) FA and the supernatant was combined with the recovered digestion mix. To remove residual beads from the peptide solution, the mix was passed over a home-made two-disc glass microfiber membrane (GE Healthcare, USA; poresize 1.2 µm, thickness 0.26 mm) tip.

**Sample clean-up for LC–MS**

Peptides were desalted on home-made $C_{18}$ StageTips[18] containing two layers of an octadecyl silica membrane (3M, USA). All centrifugation steps were carried out at room temperature. The StageTips were first activated and equilibrated by passing 50 µL of methanol (600 × *g*, 2 min), 80% (v/v) acetonitrile (ACN) with 0.5% (v/v) FA (600 × *g*, 2 min) and 0.5% (v/v) FA (800 × *g*, 3 min) over the tips. Next, the tryptic digests were passed over the tips (800 × *g*, 3-4 min). The flow-through was collected and applied a second time (same settings). The immobilized peptides were then washed with 50 µL and 25 µL 0.5% (v/v) FA (800 × *g*, 3 min). Bound peptides were eluted from the StageTips by application of two rounds of 25 µL 80% (v/v) ACN with 0.5% (v/v) FA (600 × *g*, 2 min). After elution from the StageTips, the peptide samples were dried using a vacuum concentrator (Eppendorf, Germany) and the peptides were dissolved in 15 µl 0.1% (v/v) FA prior to analysis by MS.

**LC-MS/MS analysis**

LC–MS/MS experiments were performed on an Orbitrap Fusion Lumos Tribrid instrument (Thermo Scientific, USA) that was coupled to an EASY-nLC 1200 liquid chromatography (LC) system (Thermo Scientific, USA). The LC was operated in the one-column mode. The analytical column was a fused silica capillary (75 µm × 46 cm) with an integrated PicoFrit emitter (New Objective, USA) packed in-house with Reprosil-Pur 120 C18-AQ 1.9 µm resin (Dr. Maisch, Germany). The analytical column was encased by a PRSO-V2 column oven (Sonation, Germany) and attached to a nanospray flex ion source (Thermo Scientific, USA). The column oven temperature was adjusted to 50 °C during data acquisition. The LC was equipped with two mobile phases: solvent A (0.1% (v/v) FA in water) and solvent B (0.1% (v/v) FA in 80% (v/v) ACN). All solvents were of UPLC grade (Honeywell, USA). Peptides were directly loaded onto the analytical column with a maximum flow rate that would not exceed the set pressure limit of 980 bar (usually around 0.5–0.8 µl min-1). Peptides were subsequently separated on the analytical column by running a 200 min gradient of solvent A and solvent B at a flow rate of 300 nl min-1 (gradient: start with 9% solvent B; gradient 9–40% solvent B for 180 min; gradient 40–100% solvent B for 15 min and 100% solvent B for 5 min). The mass spectrometer was operated using Xcalibur software (version 4.3.73.11; Thermo Fischer Scientific) and was set in the positive ion mode. The ionization potential (spray voltage) was set to 2.3 kV. A top-speed data-dependent method with a cycle time of 3 seconds was selected for data acquisition. Precursor ion scanning ($MS^1$) was performed in the Orbitrap analyzer (FTMS; Fourier Transform Mass Spectrometry) at a resolution of 120,000 FWHM (full width at half maximum @ 200 m/z) in the scan range of *m/z* 375–1,500 with the internal lock mass option turned on (lock mass was *m/z* 445.12002, polysiloxane)[19]. The automatic gain control (AGC) was set to “standard” and the maximum injection time was machine determined (“auto”). Product ion spectra were recorded in the Orbitrap at a resolution of 15,000 FWHM. The scan range for $MS^2$ was set to “auto”. Ions for fragmentation were selected in the quadrupole (isolation window of *m/z* 1.6) based on their intensity (threshold $5 \times 10^4$ ions) and charge state (only charge state of 2-7) in the full survey scan. The AGC target was set to “standard” and the maximum injection time to “auto”. Selected precursor ions were fragmented by Higher-energy C-trap dissociation (HCD) with normalized collision energy (NCE) set to 30%. Monoisotopic precursor selection was enabled. During $MS^2$ data acquisition, dynamic ion exclusion was set to 60 s with a repeat count of 1 and a mass tolerance of ± 10 ppm.

**Peptide and Protein identification using MaxQuant and Perseus**

RAW spectra were submitted to an Andromeda[20] search in MaxQuant (version 1.6.17.0) using the default settings[21]. Label-free quantification was activated[22]. MS/MS spectra data were searched against the self-assembled metaproteome databases of the ‘arkashin shurf’ (45 649 entries) or the ‘helicopter spring’ (99 930 entries), accordingly. All searches included a contaminants database (as implemented in MaxQuant, 246 sequences). The contaminants database contains known MS contaminants and was included to estimate the level of contamination. Andromeda searches allowed for oxidation of methionine residues (16 Da) and acetylation of the protein N-terminus (42 Da) as dynamic modifications while carbamidomethylation of cysteine residues (57 Da, alkylation with IAM) was selected as static modification. Enzyme specificity was set to “Trypsin/P”. The instrument type in Andromeda searches was set to Orbitrap and the precursor mass tolerance was set to ±20 ppm (first search) and ±4.5 ppm (main search). The MS/MS match tolerance was set to ±20 ppm.

The peptide spectrum match FDR and the protein FDR were set to 0.01 (based on target-decoy approach). Minimum peptide length was 7 amino acids. For protein quantification, unique and razor peptides were allowed. In addition to unmodified peptides, modified peptides with dynamic modifications were allowed for quantification. The minimum score for modified peptides was set to 40.

Further data analysis and filtering of the MaxQuant output was done in Perseus[23] (version 1.6.14.0). Label-free quantification (LFQ) intensities were loaded into the matrix from the proteinGroups.txt file and potential contaminants as well as reverse hits from the reverse database and hits only identified based on peptides with a modification site were removed.

Biological replicates of the unlabeled controls and the FP-alkyne labeled samples were combined into two categorical groups to allow comparison of the samples. The data were transformed to the log2-scale and only those protein groups with a minimum of 2 identified unique peptides were kept in the matrix. Furthermore, only hits with a valid LFQ intensity for at least one of the probe-labeled sample replicates were selected for further analysis. Prior to quantification, missing values were imputed from a normal distribution (width 0.3, down shift 1.8). Comparison of normalized protein group quantities (relative quantification) between different MS runs was solely based on the LFQ intensities as calculated by MaxQuant (MaxLFQ algorithm)[22]. Briefly, label-free protein quantification was switched on and unique and razor peptides were considered for quantification with a minimum ratio count of 2. Retention times were recalibrated based on the built-in nonlinear time-rescaling algorithm. MS/MS identifications were transferred between LC-MS/MS runs with the “Match between runs” option in which the match time window was set to 0.7 min and the alignment time window to 20 min. The quantification was based on the “value at maximum” of the extracted ion current. At least two quantitation events were required for a quantifiable protein. The log2-fold enrichment of protein groups with FP-alkyne was calculated based on the mean LFQ intensity compared to the DMSO control. Protein groups with a negative fold enrichment were excluded from further analysis. The remaining protein groups were reported in the respective figure (Fig. 4).

**Bioinformatic analyses of enriched proteins**

In order to confidently predict potential serine hydrolases among the group of proteins that was enriched with FP-alkyne, the protein sequences of the respective proteins were analyzed using various tools and databases. Sequence similarity to deposited serine hydrolases including the presence of characteristic serine hydrolase domains was analyzed using UniRef100[6], PFAM[24], NCBI CDD[25] and InterProScan[26]. Structural homology to known serine hydrolases was assessed using the SWISS-MODEL template library[27] and HHpred[28].

**Heterologous protein expression and purification**

The putative esterase selected for biochemical characterization (identifier: ExploCarb_3811S_S4_483_length_13114_cov_941_5) was heterologously expressed in *E. coli*. Thereto, *E. coli* Rosetta cells (Novagen, USA) were transformed with the commercially obtained construct (BioCat, Germany) of the codon-optimized gene cloned into a pET-28b(+) vector encoding a C-terminal 6× His-tag. For recombinant expression of the UPF0227 gene, a freshly inoculated 1 L culture in LB medium supplemented with 50 µg $mL^{-1}$ kanamycin and 50 µg $mL^{-1}$ chloramphenicol was grown to an $OD_{600}$ of 0.4 at 37 °C with constant shaking (180 rpm) until subsequent induction of the protein expression with 500 µM isopropyl-β-D-thiogalactopyranoside (IPTG). Upon further incubation at 18 °C for 16 h, the cells were harvested by centrifugation (8,000 × *g*, 4 °C, 20 min) and the resuspended in 5 mL 50 mM Tris-HCl pH 7.0 per gram wet weight of the pellet. Cell lysis was performed by sonication in three cycles for 5 min (cycle 0.5, amplitude 50) with a UP 200S sonicator (Hielscher Ultrasonics, Germany). The crude extract was cleared by centrifugation (12,000 × *g*, 45 min, 4 °C) and the lysate was passed through a 0.45 µm filter. Protein affinity purification was done using a Protino™ Ni-TED 1000-packed column (Macherey-Nagel, Germany) according to the manufacturer's instructions. Prior to further clean-up of the recombinant protein by size exclusion chromatography (SEC), the elution buffer was exchanged with size exclusion buffer (50 mM Tris-HCl pH 7.5, 20 mM NaCl) by centrifugation (6,000 × *g*, room temperature, 40 min) using Amicon® centrifugal filter devices (10 kDa cutoff, Merck, Germany) and the solution was concentrated to 1 mL. SEC was performed on a HiLoad® 16/600 Superdex® 200 pg column (GE Healthcare, USA) connected to an ÄKTA™ FPLC system (GE Healthcare, USA) at a flow rate of 1 mL $min^{-1}$. Fractions containing the recombinant protein (detection at 280 nm) were pooled and concentrated as described above. For long-time storage at -80 °C, 50% (v/v) glycerol was added to the protein solution that was flash-frozen in liquid nitrogen.

**Structural and bioinformatics analysis of the putative esterase**

Retrieval of homologous sequences, structures and domains was conducted with BLAST (blastp tool)[29], HHpred[28] and HMMER (phmmer tool)[30], respectively. For further structural analysis and structure comparison, a model of the UPF0227 protein was constructed with AlphaFold (version 2.0) using default settings[31]. The resulting PDB file was used for visualization and processing of the protein structure with UCSF Chimera[32] (version 1.14).

**Biochemical characterization of the heterologous protein**

The activity of the heterologously expressed UPF0227 protein was determined using a continuous assay with the chromogenic *para*-nitrophenyl (pNP) substrates pNP-acetate, pNP-butyrate, pNP-octanoate, pNP-decanoate and pNP-dodecanoate (all obtained from Megazyme, Ireland). The increase in absorbance at 384 nm was measured using a Specord 210® photometer (Analytik Jena, Germany) and the enzymatic activity was calculated from a calibration curve with pNP. The pH and temperature optimum of the enzyme was determined with pNP-butyrate prior to assessing the enzyme kinetics with the different pNP-esters at the respective pH and temperature using substrate concentrations up to 0.7 mM. To test the inhibition effect of the FP-alkyne probe used for ABPP, 6 µg/ml protein in 50 mM Tris-HCl pH 7.5, 20 mM NaCl were incubated with 2 or 4 µM of the probe for 10 min at 70 °C. The protein solutions were then submitted to activity measurement with pNP-acetate to determine the residual esterase activity compared to a DMSO-treated control.

***In vitro* labeling**

*In vitro* ABPP with FP-alkyne was performed by incubation of indicated amounts of the enzyme with 4 µM of the probe (1 h, 70 °C) in a final reaction volume of 50 µL. An equal volume of DMSO was added to the negative controls. Preincubation with paraoxon-ethyl (Sigma-Aldrich, USA), if applicable, was done at a final concentration of 100 µM (15 min, 70 °C). The subsequent click reaction was performed as described above (Click reaction and affinity purification) using Cy3-$N_3$ (synthesized in house) as click tag. For gel-based analysis, the samples were mixed with 1 equivalent 4× LDS gel loading dye (423 mM Tris HCl, 563 mM Tris base, 8% (w/v) lithium dodecyl sulfate (LDS), 40% (w/v) glycerol, 2 mM EDTA, 0.075% (w/v) SERVA Blue G250; supplemented with 100 mM DTT) and incubated at 70 °C for 15 min. Separation of proteins (1/5 of the initial amount) by gel electrophoresis was done on a 11% Bis-Tris resolving gel, followed by visualization of labeled proteins using a Typhoon FLA 9000 laser scanner (GE Healthcare, USA).

### Supplementary References


1 Gavrilov, S. N. *et al.* Isolation and Characterization of the First Xylanolytic Hyperthermophilic Euryarchaeon Thermococcus sp. Strain 2319x1 and Its Unusual Multidomain Glycosidase. *Front. Microbiol.* **7**, 552 (2016).

2 Joshi, N. & Fass, J. Sickle: a sliding-window, adaptive, quality-based trimming tool for FastQ files (version 1.33) [Software]. (2011).

3 Nurk, S., Meleshko, D., Korobeynikov, A. & Pevzner, P. A. metaSPAdes: a new versatile metagenomic assembler. *Genome Res.* **27**, 824-834 (2017).

4 Hyatt, D. *et al.* Prodigal: prokaryotic gene recognition and translation initiation site identification. *BMC Bioinformatics* **11**, 119 (2010).

5 Bornemann, T. L. V., Esser, S. P., Stach, T. L., Burg, T. & Probst, A. J. uBin – a manual refining tool for metagenomic bins designed for educational purposes. *BioRxiv* 2020.2007.2015.204776 (2020).

6 Suzek, B. E. *et al.* UniRef clusters: a comprehensive and scalable alternative for improving sequence similarity searches. *Bioinformatics* **31**, 926-932 (2015).

7 Langmead, B. & Salzberg, S. L. Fast gapped-read alignment with Bowtie 2. *Nat. Methods* **9**, 357-359 (2012).

8 Wu, Y. W., Simmons, B. A. & Singer, S. W. MaxBin 2.0: an automated binning algorithm to recover genomes from multiple metagenomic datasets. *Bioinformatics* **32**, 605-607 (2016).

9 Brown, C. T. *et al.* Unusual biology across a group comprising more than 15% of domain Bacteria. *Nature* **523**, 208-211 (2015).

10 Dick, G. J. *et al.* Community-wide analysis of microbial genome sequence signatures. *Genome Biol.* **10**, R85 (2009).

11 Sieber, C. M. K. *et al.* Recovery of genomes from metagenomes via a dereplication, aggregation and scoring strategy. *Nat. Microbiol.* **3**, 836-843 (2018).

12 Chaumeil, P. A., Mussig, A. J., Hugenholtz, P. & Parks, D. H. GTDB-Tk: a toolkit to classify genomes with the Genome Taxonomy Database. *Bioinformatics* **36**, 1925-1927 (2020).

13 Parks, D. H., Imelfort, M., Skennerton, C. T., Hugenholtz, P. & Tyson, G. W. CheckM: assessing the quality of microbial genomes recovered from isolates, single cells, and metagenomes. *Genome Res.* **25**, 1043-1055 (2015).

14 Tatusova, T. *et al.* NCBI prokaryotic genome annotation pipeline. *Nucleic Acids Res.* **44**, 6614-6624 (2016).

15 Wang, W. *et al.* Protein extraction for two-dimensional electrophoresis from olive leaf, a plant tissue containing high levels of interfering compounds. *Electrophoresis* **24**, 2369-2375 (2003).

16 Benndorf, D. *et al.* Improving protein extraction and separation methods for investigating the metaproteome of anaerobic benzene communities within sediments. *Biodegradation* **20**, 737-750 (2009).

17 Wessel, D. & Flügge, U. I. A Method for the Quantitative Recovery of Protein in Dilute-Solution in the Presence of Detergents and Lipids. *Anal. Biochem.* **138**, 141-143 (1984).

18 Rappsilber, J., Mann, M. & Ishihama, Y. Protocol for micro-purification, enrichment, pre-fractionation and storage of peptides for proteomics using StageTips. *Nat. Protoc.* **2**, 1896-1906 (2007).

19 Olsen, J. V. *et al.* Parts per million mass accuracy on an orbitrap mass spectrometer via lock mass injection into a C-trap. *Mol. Cell Proteomics* **4**, 2010-2021 (2005).

20 Cox, J. *et al.* Andromeda: A Peptide Search Engine Integrated into the MaxQuant Environment. *J. Proteome Res.* **10**, 1794-1805 (2011).

21 Cox, J. & Mann, M. MaxQuant enables high peptide identification rates, individualized p.p.b.-range mass accuracies and proteome-wide protein quantification. *Nat. Biotechnol.* **26**, 1367-1372 (2008).
22 Cox, J. *et al.* Accurate Proteome-wide Label-free Quantification by Delayed Normalization and Maximal Peptide Ratio Extraction, Termed MaxLFQ. *Mol. Cell Proteomics* **13**, 2513-2526 (2014).
23 Tyanova, S. *et al.* The Perseus computational platform for comprehensive analysis of (prote)omics data. *Nat. Methods* **13**, 731-740 (2016).
24 Mistry, J. *et al.* Pfam: The protein families database in 2021. *Nucleic Acids Res.* **49**, D412-D419 (2021).
25 Lu, S. *et al.* CDD/SPARCLE: the conserved domain database in 2020. *Nucleic Acids Res.* **48**, D265-D268 (2020).
26 Jones, P. *et al.* InterProScan 5: genome-scale protein function classification. *Bioinformatics* **30**, 1236-1240 (2014).
27 Biasini, M. *et al.* SWISS-MODEL: modelling protein tertiary and quaternary structure using evolutionary information. *Nucleic Acids Res.* **42**, W252-W258 (2014).
28 Zimmermann, L. *et al.* A Completely Reimplemented MPI Bioinformatics Toolkit with a New HHpred Server at its Core. *J. Mol. Biol.* **430**, 2237-2243 (2018).
29 McGinnis, S. & Madden, T. L. BLAST: at the core of a powerful and diverse set of sequence analysis tools. *Nucleic Acids Res.* **32**, W20-W25 (2004).
30 Potter, S. C. *et al.* HMMER web server: 2018 update. *Nucleic Acids Res.* **46**, W200-W204 (2018).
31 Jumper, J. *et al.* Highly accurate protein structure prediction with AlphaFold. *Nature* **596**, 583-589 (2021).
32 Pettersen, E. F. *et al.* UCSF chimera - A visualization system for exploratory research and analysis. *J. Comput. Chem.* **25**, 1605-1612 (2004).
